# Super-resolution microscopy reveals that disruption of ciliary transition zone architecture is a cause of Joubert syndrome

**DOI:** 10.1101/142042

**Authors:** Xiaoyu Shi, Galo Garcia, Julie C. Van De Weghe, Ryan McGorty, Gregory J. Pazour, Dan Doherty, Bo Huang, Jeremy F. Reiter

## Abstract

Diverse human ciliopathies, including nephronophthisis (NPHP), Meckel syndrome (MKS) and Joubert syndrome (JBTS), can be caused by mutations affecting components of the transition zone, a ciliary domain near its base. The transition zone controls the protein composition of the ciliary membrane, but how it does so is unclear. To better understand the transition zone and its connection to ciliopathies, we defined the arrangement of key proteins in the transition zone using two-color stochastic optical reconstruction microscopy (STORM). This mapping revealed that NPHP and MKS complex components form nested rings comprised of nine-fold doublets. The NPHP complex component RPGRIP1L forms a smaller diameter transition zone ring within the MKS complex rings. JBTS-associated mutations in *RPGRIP1L* disrupt the architecture of the MKS and NPHP rings, revealing that vertebrate RPGRIP1L has a key role in organizing transition zone architecture. JBTS-associated mutations in *TCTN2*, encoding an MKS complex component, also displace proteins of the MKS and NPHP complexes from the transition zone, revealing that RPGRIP1L and TCTN2 have interdependent roles in organizing transition zone architecture. To understand how altered transition zone architecture affects developmental signaling, we examined the localization of the Hedgehog pathway component SMO in human fibroblasts derived from JBTS-affected individuals. We found that diverse ciliary proteins, including SMO, accumulate at the transition zone in wild type cells, suggesting that the transition zone is a way station for proteins entering and exiting the cilium. JBTS-associated mutations in *RPGRIP1L* disrupt SMO accumulation at the transition zone and the ciliary localization of SMO. We propose that the disruption of transition zone architecture in JBTS leads to a failure of SMO to accumulate at the transition zone, disrupting developmental signaling in JBTS.

## INTRODUCTION

Cilia are cellular projections of organisms as diverse as green algae and mammals. Conserved across these large phylogenetic distances are a common set of structural features that define cilia: all cilia are generated from a basal body, a cylinder comprised of triplet microtubules arranged with nine-fold symmetry^1–4^. Two of the three basal body microtubules project distally to form the ciliary axoneme^5^. The axoneme is ensheathed by a ciliary membrane that is contiguous with the plasma membrane. Between the basal body and the axoneme, and separating the plasma membrane from the ciliary membrane, is the transition zone. Electron micrographs of the transition zone from diverse organisms have identified Y-shaped densities, called Y links, with the stem anchored at the microtubule doublet and the two arms attached to the ciliary membrane^6^. The set of molecular components that comprise the Y links has been unclear, as has the arrangement of transition zone proteins.

Ciliopathies are human diseases caused by defective ciliary function. Several ciliopathies are caused by mutations in genes encoding transition zone components^7,8^. *NPHP1, NPHP4, and RPGRIP1L* encode transition zone components and are mutated in NPHP, characterized by kidney cysts and fibrosis^9,10^. Another subset of transition zone genes including, *B9D1, TCTN1, TCTN2, TMEM231*, and *RPGRIP1L*, are mutated in MKS, a lethal ciliopathy characterized by kidney cysts, polydactyly and occipital encephalocele. Mutations in *NPHP1, TCTN1, TCTN2, TMEM231, RPGRIP1L* and *ARL13B* also cause JBTS, characterized by hypoplasia of the cerebellar vermis^11–18^. JBTS-affected individuals exhibit hypotonia, ataxia, and cognitive disability, and can exhibit kidney cysts, polydactyly, liver fibrosis, and retinal dystrophy leading to blindness^19^. Consistent with their associations with different diseases, RPGRIP1L, NPHP1 and NPHP4 are components of a protein complex called the NPHP complex, whereas B9D1, TCTN1, TCTN2 and TMEM231 are components of a distinct complex called the MKS complex^14,20,21^. In *C. elegans*, a homolog of RPGRIP1L is required for the transition zone localization of both the NPHP and MKS complexes^22,23^. Although the means by which cilia transduce intercellular signals are becoming clearer, how JBTS-associated mutations affect those functions remains obscure.

Befitting its position at the border between the plasma and ciliary membranes, the transition zone helps define the composition of the ciliary membrane. Genetic disruption of the transition zone in the mouse, *C. elegans*, or *C. reinhardtii* can disrupt the ciliary localization of membrane-associated proteins^24–26^. For example, an essential mediator of the vertebrate Hedgehog signaling pathway, Smoothened (SMO), and a regulator of kidney epithelial growth, Polycystic Kidney disease 2 (PKD2), require the transition zone proteins B9D1, TMEM231 and TCTN2 to accumulate within the ciliary membrane^14–16,21^. Consequently, loss of any of several transition zone components leads to Hedgehog-associated developmental defects, such as polydactyly, and PKD2-associated defects, such as kidney cysts^15,16,20^.

Due to this critical role in regulating the localization of membrane proteins to the cilium, the transition zone has been proposed to act as the gate to the cilium^27^. To understand how the transition zone functions as a gate, we need to understand how it is structured at the supra-molecular, sub-organellar level. The ultrastructural features of cilia and centrosomes range in size from 200 to 500 nm. Conventional light microscopy, limited in spatial resolution by the diffraction of light, and for electron microscopy, limited in its ability to define the relative position of multiple proteins, are not equal to this task. However, super-resolution optical microscopy is well suited to resolve structures of these sizes^28–31^. For example, structured illumination microscopy (SIM)^32^ and stochastic optical reconstruction microscopy (STORM)^33^ of *Drosophila* centrosomes revealed that the centrosomal protein Pericentrin-like Protein (PLP) is organized into oriented clusters^28^. Similarly, stimulated emission depletion (STED) microscopy of the transition zone has shown that the MKS complex is ~150 nm distal to the CENTRIN edge along the ciliary axis in retinal pigmented epithelial (RPE1) cells^30^.

To elucidate the architecture of the ciliary gate, we combined multicolor three dimensional (3D) SIM, two-color 3D STORM and quantitative image analysis^34,35^. Using this approach, we found that the ciliopathy-associated proteins RPGRIP1L, TMEM231, NPHP1, and B9D1 each form a ring of a unique diameter that lies outside the microtubule axoneme close to or at the ciliary membrane. These rings are not continuous as would be expected for a membrane diffusion barrier, but are comprised of discrete clusters spaced at non-uniform but non-random distances. Together with proximodistal measurements along the ciliary axis, we used diameter and radial distribution measurements to construct a three dimensional map of the transition zone at a spatial resolution of 15–30 nm, revealing that the location of the MKS and NPHP complexes is consistent with their localization to Y links. Analysis of this transition zone architecture in human cells derived from JBTS-affected individuals indicated that RPGRIP1L and TCTN2 are essential for the localization of both the NPHP complex and the MKS complex. Recent single molecule studies revealed that SMO pauses at multiple binding sites at an area near the ciliary base^36^. Our two-color STORM analysis revealed that SMO accumulates as a ring of discrete clusters specifically in the transition zone, suggesting that the transition zone is a point at which proteins like SMO pause on their way to or from the ciliary membrane. We found that human JBTS-associated *RPGRIP1L* mutations reduce the localization of SMO at the transition zone and the ciliary membrane, underscoring the importance of an intact transition zone for Hedgehog signaling in development and JBTS. The approach to determining the relative three dimensional positions of proteins used here elucidates how ciliopathy mutations disrupt transition zone architecture and will be applicable to elucidating how other human disease mutations compromise other supra-molecular, sub-organellar assemblies.

## RESULTS

### NPHP and MKS complex components form discontinuous, concentric, nested rings within the transition zone

To assess differences in the arrangement of proteins associated with the ciliopathies Meckel Syndrome, Joubert syndrome, or nephronophthisis, we began by visualizing the transition zone of mouse tracheal epithelial cells (mTECs) using multicolor 3D-SIM. mTECs produce hundreds of cilia per cell, uniformly extend their cilia from the apical surface, and orient their cilia along the plane of the epithelial surface, allowing us to correlate cilia within an image. SIM revealed that mammalian NPHP1 was distributed in a ring in the transition zone of each cilium (Figure 1A).

**Figure 1.**
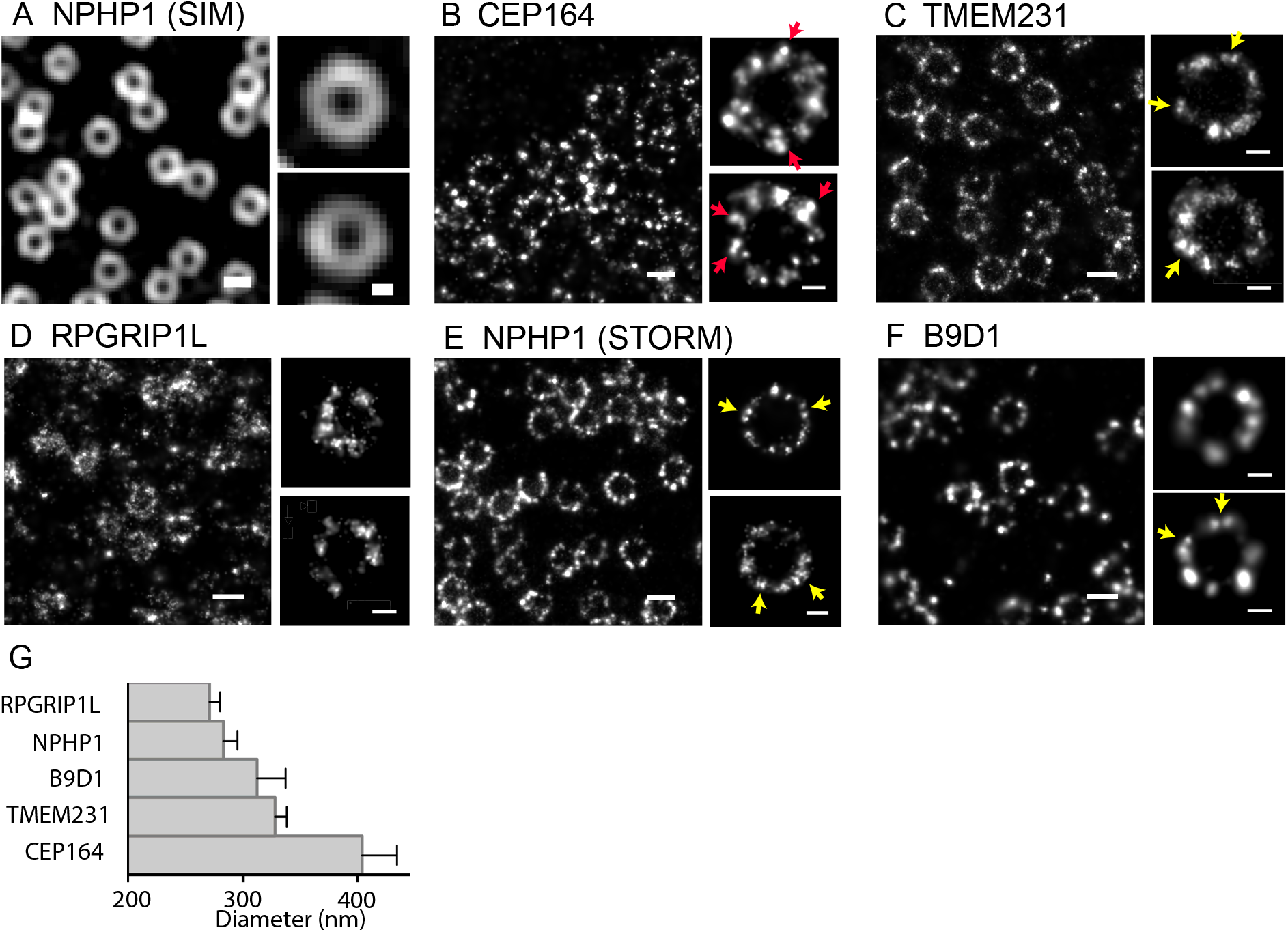
MKS and NPHP complexes form rings of discrete puncta. (**A**) SIM detection of NPHP1 in mTECs reveals a ring structure. STORM detection of (**B**) CEP164, (**C**) TMEM231, (**D**) RPGRIP1L, (**E**) NPHP1, and (**F**) B9D1 in mTECs reveals that the rings are comprised of discrete puncta. MKS complex components and NPHP1 are present in doublets oriented tangentially to the ring circumference (yellow arrows). In contrast, STORM imaging of CEP164, a distal appendage component, reveals clusters oriented obliquely to the ring circumference (red arrows). (**G**) Diameter measurements of rings in STORM images. Error bars are standard deviations. P < 0.0001 for any pair of values. Scale bars are 300 nm and 100 nm for multiple-structure images and single structure images, respectively.

The transition zone has been proposed to regulate ciliary membrane composition by acting as a membrane diffusion barrier^21^. We used single-color STORM to better resolve the distributions of ciliary proteins. The distal appendage protein CEP164 formed clusters arranged in a ring (Figure 1B), consistent with previously published observations^29^. Interestingly, we found that the rings of the transition zone proteins RPGRIP1L, NPHP1, TMEM231, and B9D1 were also comprised of discrete clusters, in contrast to continuous membrane diffusion barriers such as those based on Septin assemblies (Figure 1C-F).

Positioning of a transition zone protein in a cylindrical coordinate system (with the *z* axis along the cilium axis) requires the determination of three coordinates: the radial distance to the cilium axis, the angular position in the plane perpendicular to the cilium axis, and the proximodistal position along the cilium axis. For the radial distance, we measured the diameters of the rings formed by transition zone components by performing a robust fit^37^ of each ring structure to an ellipse (see Methods for details). Of the transition zone proteins assessed, the TMEM231 ring had the largest average diameter (328 nm ± 10 nm, *n* = 30, mean ± standard deviation shown here and for all subsequent diameter measurement results, Figure 1C and 1G). TMEM231 is a two-pass transmembrane protein likely to be present within the transition zone membrane^16^. RPGRIP1L, a protein with coiled-coil domains and C2 domains, comprised a ring of the smallest diameter (271 nm ± 9 nm, *n* = 23, Figure 1D and 1G). NPHP1 (283 nm ± 12 nm, *n* = 46) and B9D1 (312 nm ± 25 nm, *n* = 27) rings exhibited diameters intermediate to those of TMEM231 and RPGRIP1L (Figure 1E, 1F, and 1G). The distal appendage component CEP164 comprised a ring larger than that of any of the transition zone components (404 nm ± 30 nm, *n* = 31) (Figure 1B and 1G). These measurements suggest that these transition zone proteins comprise concentric nested rings, with CEP164 outside the transition zone, TMEM231 coincident with the transition zone membrane and B9D1, NPHP1, and RPGRIP1L rings being successively closer to the axoneme (Figure 1G).

### The NPHP and MKS complexes are organized into nine-fold symmetric doublets

As the transition zone rings were comprised of non-continuous clusters, we investigated whether these clusters were randomly or non-randomly positioned along the ring circumference. Because of the imperfect labeling efficiency of primary and secondary antibodies, we often observed incomplete rings with potentially missing clusters. For example, the distal appendage component CEP164 typically showed 6 to 9 clusters (Figure 1B), similar to previously observations using STED microscopy^29^. To compensate for imperfect labeling, we aligned and averaged the STORM images of multiple structures using an algorithm similar to that used for averaging single-particle cryo-electron microscopy images (see Methods for details). The averaged image of CEP164 clearly showed 9 clusters with a symmetrical distribution around the ring (Figure 2A, right). In striking contrast, many TMEM231 and NPHP1 rings contained more than 9 clusters (Figures 1C and E), some of which were doublets oriented tangential to the ring circumference (yellow arrows in Figure 1C, E and F). Also unlike CEP164, the averaged images of transition zone proteins NPHP1 and TMEM231 did not reveal nine-fold symmetry (Figures 2B Right, C Right and D Right), suggesting that the arrangement of transition zone proteins is more complex or heterogeneous than that of distal appendages. Moreover, increasing the number of images for averaging did not improve the result, further implicating structural heterogeneity.

**Figure 2.**
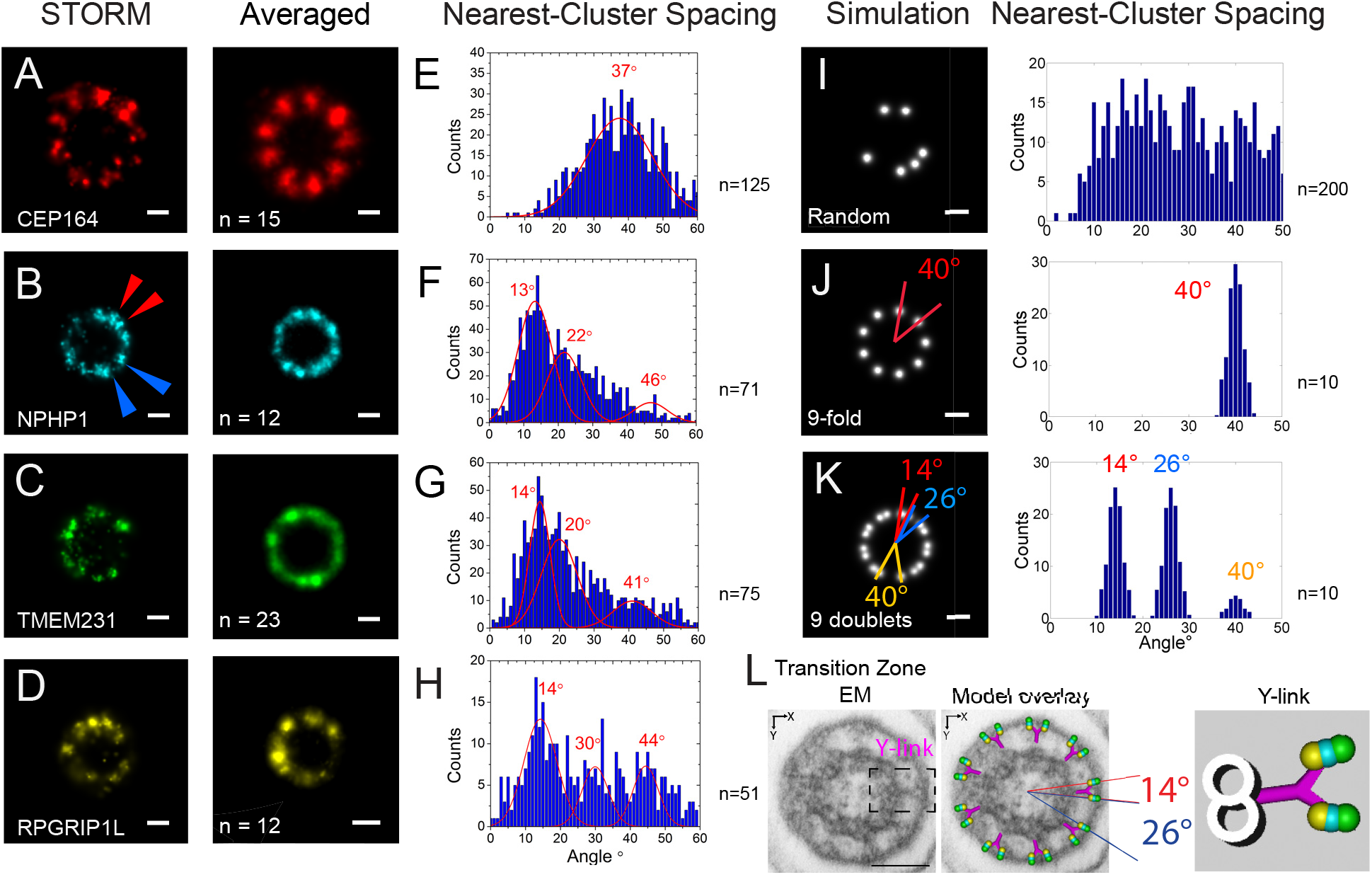
MKS and NPHP complex rings are comprised of doublets. **A-D**: STORM images of mTEC cilia stained for (**A**) CEP164, (**B**) NPHP1, (**C**) TMEM231, or (**D**) RPGRIP1L (left). Multiple images were aligned by cross correlation and averaged (right). n represents number of STORM images used for the averaging. E-H: Histograms of the angular spacing between the nearest clusters of (**E**) CEP164, (**F**) NPHP1, (**G**) TMEM231, and (**H**) RPGRIP1L rings. n represents number of STORM images used for the analysis. **I-K**: Histograms of the angular spacing of between the nearest clusters of simulated rings in which the puncta are (**I**) randomly distributed along the ring circumference, (**J**) nine-fold symmetric, and (**K**) comprised of nine-fold symmetric doublets. n represents number of simulated rings used for the analysis. (**L, left**) Transmission electron micrograph (TEM) of a transverse section through the transition zone of a mouse ependymal cell cilium. A Y link is highlighted by a dashed line. (**L, center**) Overlay of a model of angular and radial arrangement of transition zone proteins with the TEM. (**L, right**) A model of the radial and angular distributions of transition zone proteins superimposed onto a Y link structure. Scale bars, 100 nm.

To analyze the angular relationship between these clusters more quantitatively, we generated an angular position histogram of STORM localization points for each ring structure, identified clusters in the histogram using multi-Gaussian-peak fitting, calculated the spacing between the center of adjacent clusters, and then pooled nearest-cluster spacing values from all images to generate a final histogram for each analyzed protein (Figures 2E-H) (see Methods for details). When focusing on the smaller angle range, this nearest-cluster spacing analysis was less sensitive to continuous structural deformations than is global image alignment and averaging. Unlabeled clusters generated nearest-cluster spacing values in a larger angle range that are not included in subsequent analyses. In nearest-cluster spacing analysis, clusters of random angular distribution resulted in no detectable peaks (Figure 2I), whereas 9 clusters symmetrically distributed around a ring resulted in a peak at 40° (Figure 2J). The latter was close to the observed angle for CEP164 (Figure 2E) (peak position at 37 ± 2°, *n* = 125 structures, uncertainty calculated by bootstrapping here and for subsequent angular analyses).

Consistent with the observation of more than nine NPHP1, TMEM231 and RPGRP1L clusters per ring, the spacing between most clusters of these proteins was smaller than 40° (Figures 2F-H). Multi-Gaussian-peak fitting results of the histograms consistently converged to three peaks (> 80% of the boot-strapped histograms producing 3-peak fitting results even in the noisiest case of RPGRIP1L), two major peaks at around 14° and 23° (13 ± 1° and 23 ± 2° for NPHP1, *n* = 71 structures, 14 ± 1° and 20 ± 1° for TMEM231, *n* = 75 structures, and 14 ± 1° and 30 ± 1° for RPGRIP1L, *n* = 51 structures), and a minor peak at around 40°. This distribution was analogous to that produced by a simulated ring consisting of 9 symmetric doublets of clusters with one cluster unlabeled (Figure 2K). This simulation generated two major peaks, corresponding to intra-doublet and inter-doublet angles, which sum to 40°. In addition, the simulation generated a third, minor peak at 40°, corresponding to inter-doublet angles in which one cluster is unlabeled. Thus, the arrangement of rings composed by individual MKS and NPHP complex proteins can be modeled as a planar wheel comprised of nested rings comprised of nine symmetrically distributed doublets, each having an intra-doublet spacing of about 14°.

The observed doublet arrangement is strikingly similar to the ultrastructural features observed in electron micrographs of the transition zone of mouse ependymal cells (Figure 2L). TEMs show densities spanning the microtubule doublets and the ciliary membrane that form the shape of the letter Y, called Y links (e. g., Figure 2L inside of dashed outline). The angle between the tips of Y links (intra-Y link) of mouse ependymal cell electron micrographs is 12° − 25° (average = 20°, standard deviation = 2.3°, n = 70). Superimposing the transition zone assembly based on STORM measurements onto the TEM images further highlights the similarity between the MKS and NPHP complex localizations and the tips of the Y links (Figure 2L).

### NPHP1 and the MKS complex occupy the same proximodistal position in the transition zone

To map the third axis required to build a full model of the transition zone, we measured the axial positions of the transition zone rings along the ciliary axis. To register multiple transition zone components with the CEP164 ring as a common reference point, we used two-color 3D STORM. More specifically, we devised a self-aligned two-color STORM scheme using two-channel imaging of two spectrally overlapping reporter dyes to avoid the need for a separate channel registration (see Methods for details). We performed our measurements using both motile mTEC cilia and IMCD3 primary cilia. Two-color 3D-STORM confirmed that the CEP164 ring is larger than the NPHP1 ring and revealed that the two rings are concentric (Figure 3A). Using side views, we fitted the axial projections of CEP164 and NPHP1 with two Gaussian peaks, which revealed that the NPHP1 ring is 103 nm ± 36 nm (mean ± standard deviations here and for subsequent distance and thickness measurements, *n* = 5) distal to the CEP164 ring and has a thickness of 122 ± 36 nm (width at half maximum of the Gaussian fitting).

**Figure 3.**
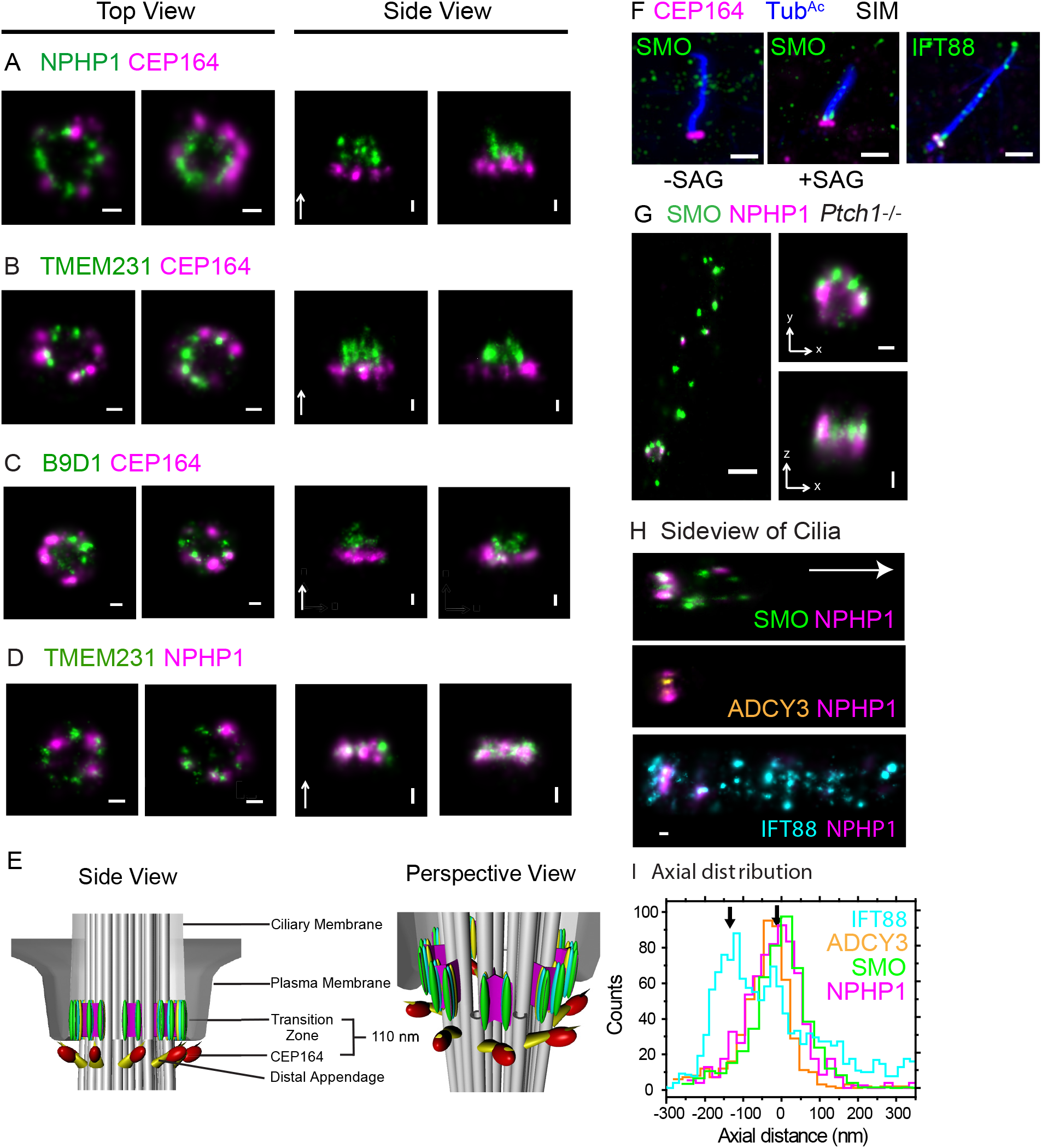
MKS and NPHP complexes occupy the same proximodistal position in the transition zone, where SMO docks upon Hh pathway activation. (**A-C**) Two-color STORM images of CEP164 (magenta) and (**A**) NPHP1 (green), (**B**) TMEM231 (green), and (**C**) B9D1 (green). Top views (left) are of mTECs (top views) and side views are of IMCD3 cells. The arrows in the side views point in the direction of the ciliary tip. Scale bars, 100 nm. (**D**) Two-color STORM images of TMEM231 (green) and NPHP1 (magenta). Scale bars, 100 nm. (**E**) Side view and perspective view of a 3D model of the ciliary gate including the microtubules of the basal body and axoneme (gray), CEP164 along the distal appendages (yellow), and TMEM231 (green), NPHP1 (teal) and RPGRIP1L (yellow) within the transition zone. (**F, left & middle**) SIM image of human fibroblasts stained for SMO (green), CEP164 (magenta) and acetylated tubulin (blue) treated with or without SAG. Scale bar, 1 μm. (**F, right**) SIM image of human fibroblasts stained for IFT88 (green), CEP164 (magenta) and acetylated tubulin (blue). Scale bar, 1 μm. (**G**) Two-color STORM of SMO (green) and NPHP1 (magenta) in *Ptch1^−/−^* MEFs. Scale bars, 500 nm (left), 100 nm (right). (**H**) Two-color STORM of side views of primary cilia. SMO (green) and NPHP1 (magenta) were imaged in *Ptch1^−/−^* MEFs. ADCY3 (orange) and NPHP1 (magenta) and were imaged in WT MEFs. The arrow points in the direction of the ciliary tip. Scale bar, 100 nm. (**I**) Histogram of the intensity of SMO (green), ADCY3 (orange), NPHP1 (magenta), and IFT88 (cyan) along the axonemal direction, corresponding to images of which (**H**) is representative.

To assess the position of the MKS complex, we also examined the axial position of the MKS complex components TMEM231 and B9D1. Two-color 3D STORM of CEP164 and either TMEM231 or B9D1 revealed that, like NPHP1, TMEM231 and B9D1 comprise rings within the concentric CEP164 ring (Figure 3B and C). Side views showed that the TMEM231 and B9D1 rings are 115 ± 16 nm (*n* = 12) and 91 ± 40 nm (*n* = 5) distal to CEP164, respectively (Figure 3B and C). The thickness of TMEM231 and B9D1 rings are 114 ± 61 and 97 ± 50 nm respectively. The similar distances of the NPHP complex member NPHP1 and the MKS complex member TMEM231 from CEP164 suggested that they occur at the same region along the ciliary proximodistal axis. Indeed, two-color 3D STORM of NPHP1 and TMEM231 confirmed that they form similar sized rings at the same proximodistal position along the cilium (Figure 3D).

With the measured radial, angular and axial distributions of transition zone components, we rendered a 3D model of protein localization within the transition zone (Figure 3E). This model illustrates that the MKS and NPHP complexes are arranged as doublets with different radial distances within the transition zone. These doublets are nine-fold symmetric, and likely to represent the arms of the Y links connecting the ciliary membrane to the Y link stem anchored at the microtubule doublet. As revealed by two color 3D STORM, the middle of the transition zone MKS and NPHP rings is approximately 110 nm distal to the distal appendages, and the transition zone extends for about 120 nm along the ciliary axis.

### SMO and other ciliary proteins accumulate at the transition zone

A critical role for the transition zone is to control the composition of the ciliary membrane. One ciliary membrane component essential for Hedgehog signaling and misactivated in diverse cancers is SMO, a seven-pass transmembrane protein^38^. SMO accumulation in the cilium and subsequent pathway activation requires the transition zone MKS complex^14–16,21^. Hedgehog binding to its receptor, PTCH1, promotes the exit of PTCH1 from the cilium, and allows SMO to accumulate in the cilium^39,40^. Loss of PTCH1 leads to constitutive SMO localization to the cilium (as observed in *Ptch1*^−/−^ mouse embryonic fibroblasts [MEFs], Supplementary Figure 1) and constitutive pathway activation, an event that causes both basal cell carcinoma^41,42^ and medulloblastoma^43,44^. Recent single molecule studies of SMO have revealed that SMO pauses at an area near the ciliary base^36^. To determine where SMO localizes at the ciliary base, we visualized the position of SMO relative to the distal appendages and the transition zone using two-color SIM and 3D STORM. Wild-type unstimulated fibroblasts do not localize SMO to the ciliary membrane, as expected (Figure 3F). Wild-type fibroblasts treated with Smoothened Agonist (SAG), a chemical activator of Hh signaling^45^, showed an accumulation of SMO at a region of the ciliary base distal to the distal appendages marked with CEP164 (Figure 3F). Similarly, *Ptch1^−/−^* MEFs, in which Hh signaling is active, displayed SMO along the cilium in addition to an accumulation near the base (Figure 3G, compared to wild type scenario in Supplementary Figure S1).

To precisely determine where SMO localizes at the ciliary base, we examined the distribution of SMO and NPHP1 using two color 3D STORM and found that SMO largely colocalizes with NPHP1 (Figure 3G). Indeed, quantitation of SMO position along the ciliary axis shows it at the same position as NPHP1 (Figure 3I), indicating that SMO accumulates at the transition zone.

Another protein that localizes to the ciliary membrane is ADCY3^46,47^, a transmembrane adenylyl cyclase involved in olfaction, adiposity, and male fertility^48–50^. Like SMO, ADCY3 accumulated at the transition zone (Figures 3H and I), suggesting that pausing at the transition zone may be a general characteristic of ciliary membrane proteins.

For comparison, we also examined the localization of intraflagellar transport (IFT)-B complex component IFT88. The IFT-B complex is required to carry many ciliary proteins across the transition zone to the ciliary tip. Consistent with previous indications that the IFT-B complex is assembled at the distal appendages^51^, IFT88 prominently colocalized with CEP164 (Figure 3F). A less prominent accumulation of IFT88 was also observed distal to the distal appendages (Figure 3F). Two color 3D STORM confirmed that IFT88 predominantly localizes 110 nm proximal to the transition zone, consistent with the distal appendages, with a smaller accumulation colocalizing with NPHP1 at the transition zone (Figures 3H and I). The enrichment of SMO, ADCY3 and IFT88 at the transition zone suggests that the transition zone is a waypoint for diverse proteins entering or leaving the cilium.

### JBTS-associated mutations in *TCTN2* disrupt MKS and NPHP complex localization to transition zone

Many of the more than 30 proteins associated with JBTS are components of the transition zone^27^. Mutations in *TCTN2* cause JBTS24^20^, but understanding of how remains obscure. To assess how TCTN2 functions in the human transition zone, we examined how JBTS-associated compound heterozygous mutations in *TCTN2* affected the composition of the transition zone and ciliary membrane. The JBTS-affected individual is compound heterozygous for a missense mutation (G373R) and a frame shift mutation (D267fsX26) in *TCTN2*, whereas the two unaffected siblings (identical twins) are heterozygous for the missense mutation (G373R) and a wild type allele^52^.

SIM and fluorescence intensity quantitation of control and *TCTN2* mutant fibroblasts revealed that TCTN2 was reduced at the transition zone by 72% ± 3% (mean difference ± standard error of difference reported here and for subsequent fluorescence intensity measurements. n = [144–4, 144–3; 82, 82]) in the *TCTN2* mutant cells (Figure 4A). To begin to examine the consequences of loss of TCTN2 function, we assessed whether TCTN2 participates in the localization of TMEM231, another MKS complex protein underlying JBTS^11^. *TCTN2* mutant fibroblasts displayed a 75% ± 5% (n = [144–4, 144–3; 90, 86]) reduction of TMEM231 at the transition zone (Figure 4A). Thus, human TCTN2 is required for the transition zone localization of TMEM231.

We further investigated whether *TCTN2* mutations also affect the NPHP complex by assessing the localization of RPGRIP1L and NPHP1, two NPHP complex members mutated in JBTS^17,53^. *TCTN2* mutant fibroblasts displayed a 53% ± 6% (n = [144–4, 144–3; 74, 50]) reduction of RPGRIP1L and 36% ± 4% (n = [144–4, 144–3; 100, 87]) reduction of NPHP1 at the transition zone (Figure 4A). Thus, human TCTN2 is required to localize wild type amounts of both NPHP and MKS complex components at the transition zone.

**Figure 4.**
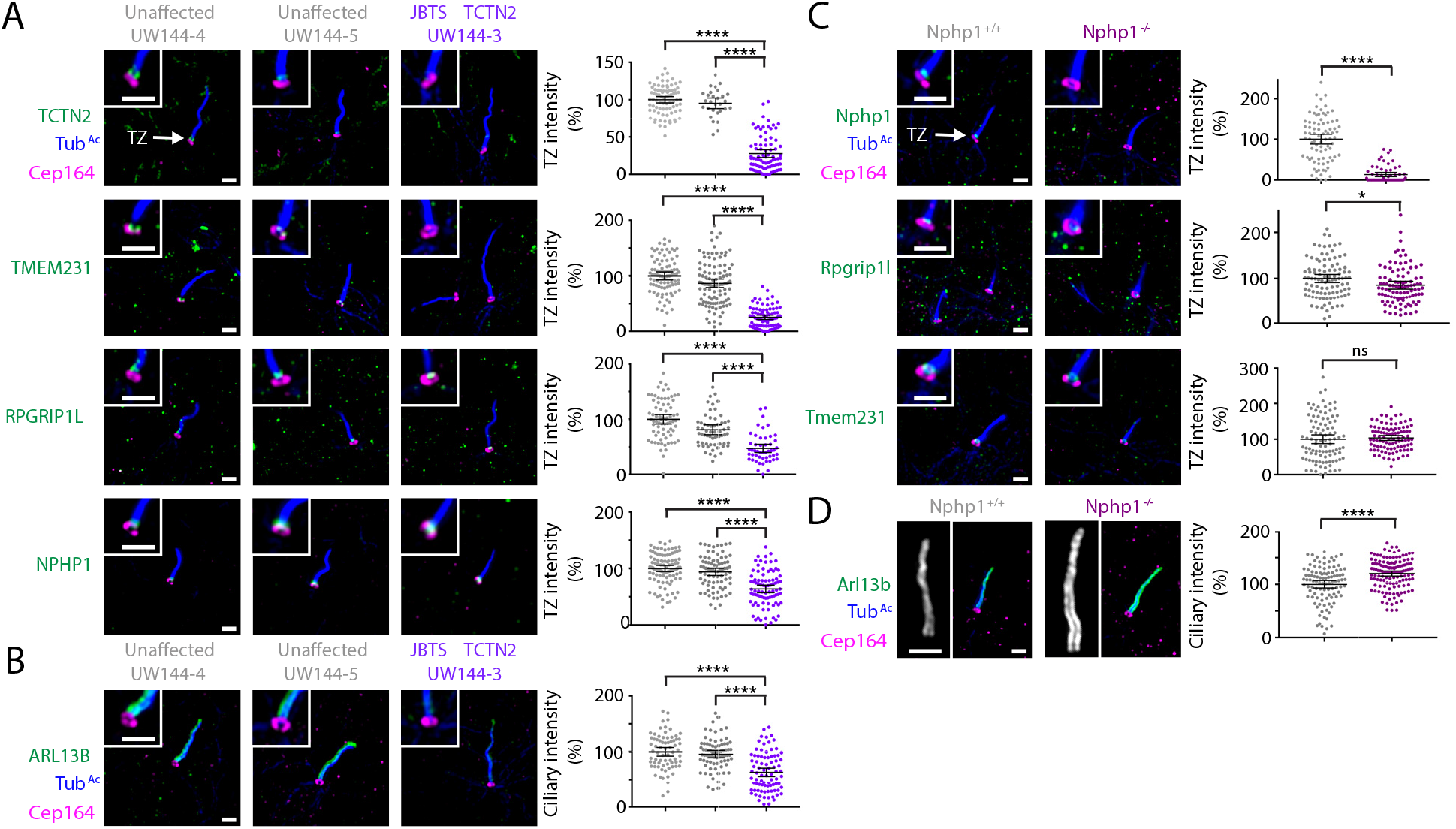
MKS and NPHP complexes are displaced from the transition zone in a Joubert syndrome patient with mutations in *TCTN2*. (**A**) SIM images of acetylated tubulin (Tub^Ac^, blue), CEP164 (magenta) and TCTN2, TMEM231, RPGRIP1L, or NPHP1 (green) in fibroblasts from a JBTS-affected individual and two unaffected siblings. Quantification of TCTN2, TMEM231, RPGRIP1L, and NPHP1 fluorescence intensity at the transition zone (right). Mean ± 95% confidence interval are pooled data from two independent experiments. n = number of transition zones measured (144–4, 144–5, 144–3): TCTN2 (82, 29, 82), TMEM231 (90, 105, 86), RPGRIP1L (74, 62, 50), and NPHP1 (100, 82, 87). (**B**) SIM images of acetylated tubulin (blue), CEP164 (magenta) and ARL13B (green). Quantification of ciliary ARL13B fluorescence intensity (right). Mean ± 95% confidence interval are pooled data from two independent experiments. n = number of cilia measured (144–4, 144–5, 144–3; 72, 78, 84). (**C**) SIM images of Tub^Ac^ (blue), CEP164 (magenta) and NPHP1, RPGRIP1L, and TMEM231 (green) in *Nphp1*^+/+^ and *Nphp1^−/−^* mouse embryonic fibroblasts. Quantification of Nphp1, Rpgrip1l, and Tmem231 fluorescence intensity at the transition zone (right). Mean ± 95% confidence interval are pooled data from three independent experiments. n = number of transition zones measured *(Nphp1*^+/+^, *Nphp1^−/−^*): NPHP1 (83, 59), RPGRIP1L (102, 102), and TMEM231 (100, 98). (D) SIM images of Tub^Ac^ (blue), CEP164 (magenta) and ARL13B (green). Quantification of ciliary Arl13b fluorescence intensity (right). Mean ± 95% confidence interval are pooled data from three independent experiments. n = number of cilia measured (*Nphp1*^+/+^, *Nphp1^−/−^*; 105, 133). ns, not significant. *, P value = 0.021. ****, P value <0.0001. Multiple comparisons were performed using an ordinary one-way ANOVA, along with Tukey’s multiple comparisons test to analyze pairwise differences. Two sample comparisons were performed using a two tailed unpaired t test. Scale bars, 1 μm.

In mice, the MKS complex is critical for the ciliary localization of ARL13B, a small GTPase^13^. As mutations in *ARL13B* also cause JBTS^54^, we examined whether JBTS-associated mutations in *TCTN2* affect the ciliary localization of ARL13B. *TCTN2* mutant fibroblasts displayed a 37% ± 5% (n = [144–4, 144–3; 72, 84]) reduction of ARL13B at the cilium (Figure 4B). Thus, JBTS-associated mutations in *TCTN2* disrupt the composition of the transition zone and diminish the ciliary localization of the JBTS-associated protein ARL13B.

### NPHP1 has a subtle role in organizing the transition zone

To assess how NPHP1 functions in the transition zone, we examined how mutation of *Nphp1* in mouse embryonic fibroblasts affected the composition of the transition zone and ciliary membrane. SIM of *Nphp1*^+/+^ control and *Nphp1^−/−^* fibroblasts revealed that, as expected, NPHP1 was absent from the transition zone of *Nphp1^−/−^* cells (100% ± 8%, n = [*Nphp1*^+/+^, *Nphp1^−/−^*; 83, 59], Figure 4C). To examine the consequences of loss of NPHP1 function, we assessed whether NPHP1 participates in the localization of RPGRIP1L. We found that RPGRIP1L was reduced by 15% ± 6% (n = [*Nphp1*^+/+^, *Nphp1^−/−^;* 102, 102]) from the transition zones of *Nphp1^−/−^* cells (Figure 4C). Thus, NPHP1 has a minor role in the transition zone localization of RPGRIP1L.

We further investigated whether NPHP1 organizes the MKS complex by assessing the localization of TMEM231. The quantity of TMEM231 in the transition zone was unaffected in *Nphp1^−/−^* cells (4% ± 7%, n = [*Nphp1*^+/+^, *Nphp1^−/−^*; 100, 98]), Figure 4C). Thus, NPHP1 is not required for the transition zone localization of a key MKS complex component.

We also examined whether NPHP1 participates in the ciliary localization of ARL13B. Surprisingly, ARL13B levels were elevated by 20% ± 4% (n = [
*Nphp1*^+/+^, *Nphp1^−/−^*; 105, 133]) in *Nphp1^−/−^* cells (Figure 4D). Thus, whereas NPHP1 has a minor role in the localization of another NPHP component, RPGRIP1L, to the transition zone, it is dispensable for localization of one MKS component and dispensable for the MKS complex-dependent localization of ARL13B to the cilium.

### JBTS mutations in *RPGRIP1L* displace MKS and NPHP complex proteins from the transition zone

To assess how RPGRIP1L functions in the human transition zone, we examined how JBTS-associated compound heterozygous mutations of three individuals from two different families affected the composition of the transition zone and ciliary membrane. The JBTS-affected individuals have missense mutations affecting the C2-N domain of RPGRIP1L in combination with a nonsense mutation (Supplementary Figure S2)^52^. SIM and fluorescence intensity quantitation of control fibroblasts from an unaffected sibling and three affected individuals revealed that the JBTS-associated mutations in *RPGRIP1L* reduced the level of RPGRIP1L at the transition zone by 81% ± 3% (n = [15–6, 15–4; 100, 65], Figure 5A). To begin to examine the consequences of loss of RPGRIP1L function, we assessed whether RPGRIP1L participates in the localization of NPHP1. We found that NPHP1 was dramatically reduced by 85% ± 7% (n = [15–6, 15–4; 101, 58]) from the transition zones of cells from JBTS-associated *RPGRIP1L* mutations (Figure 5A). Thus, human RPGRIP1L is required for the transition zone localization of NPHP1, but NPHP1 has only a minor role in localizing RPGRIP1L to the transition zone (Figure 4C).

**Figure 5.**
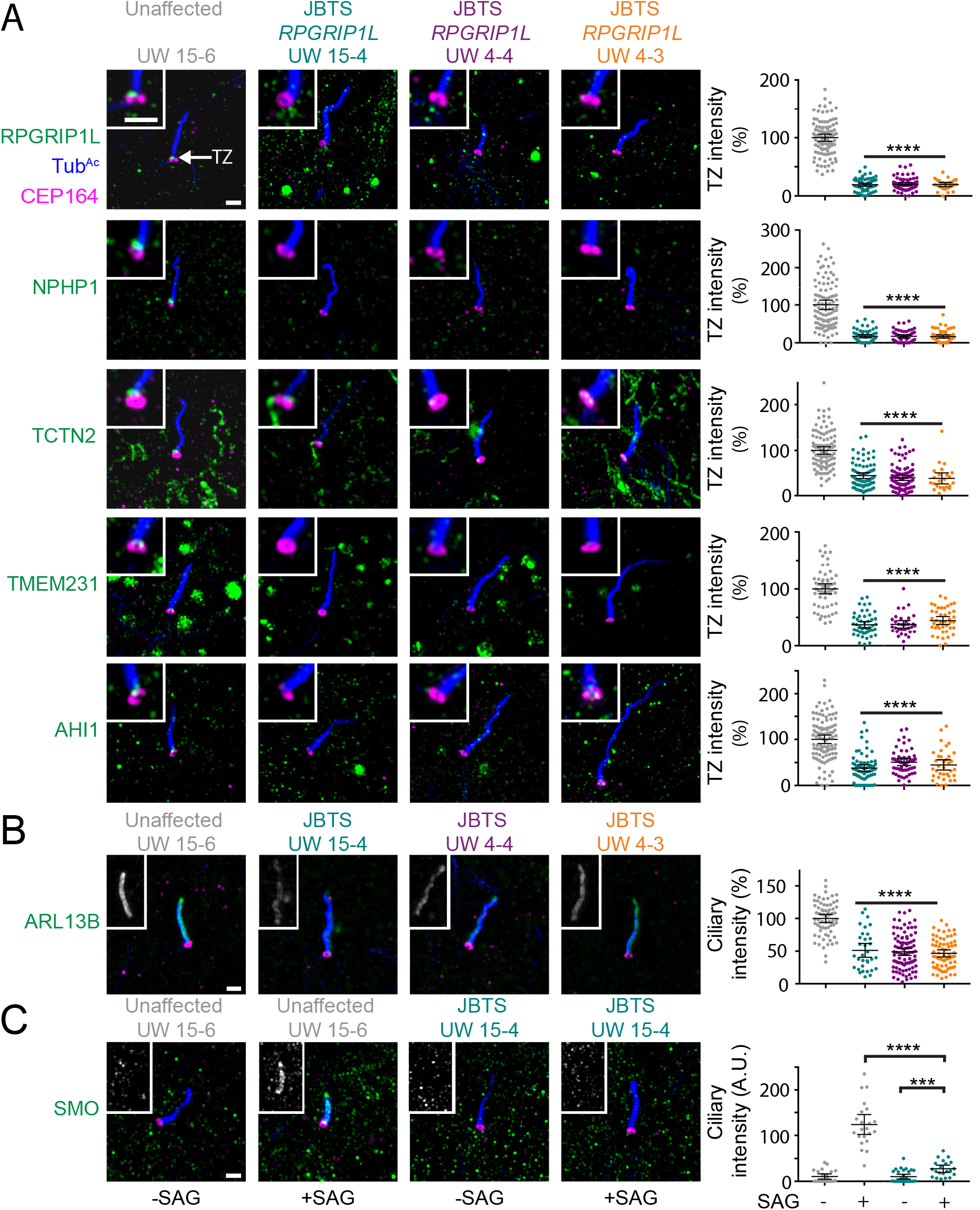
MKS and NPHP complexes are displaced from the transition zone in Joubert syndrome patients with mutations in *RPGRIP1L*. (**A**) SIM images of acetylated tubulin (Tub^Ac^, blue), CEP164 (magenta) and RPGRIP1L, NPHP1, TCTN2, TMEM231 or AHI1 (green) in fibroblasts from three JBTS-affected individuals and an unaffected sibling. Quantification of RPGRIP1L, NPHP1, TCTN2, TMEM231 and AHI1 fluorescence intensity at the transition zone (right). Mean ± 95% confidence interval are pooled data from three independent experiments. n = number of transition zones measured (15–6, 15–4, 4–4, 4–3): RPGRIP1L (100, 65, 62, 28), NPHP1 (101, 58, 53, 53), TCTN2 (102, 80, 92, 24), TMEM231 (58, 50, 33, 43), and AHI1 (100, 69, 55, 36). (**B**) SIM images of Tub^Ac^ (blue), CEP164 (magenta) and ARL13B (green). Quantification of ciliary ARL13B fluorescence intensity (right). Mean ± 95% confidence interval are pooled data from three independent experiments. n = number of cilia measured (15–6, 15–4, 4–4, 4–3; 67, 33, 94, 68). (**C**) SIM images of Tub^Ac^ (blue), CEP164 (magenta) and SMO (green) in fibroblasts from a JBTS-affected individual and an unaffected sibling in the presence or absence of SAG. Quantification of ciliary SMO fluorescence intensity (right). Mean ± 95% confidence interval is shown in graph. Two independent experiments were performed. n = number of cilia measured (15–6 no SAG, 15–6 SAG, 15–4 no SAG, 15–4 SAG; 23, 22, 29, 21). ***, P value = 0.0003. ****, P value < 0.0001. Multiple comparisons were performed using an ordinary one-way ANOVA, along with Tukey’s multiple comparisons test to analyze pairwise differences. Two sample comparisons were performed using a two tailed unpaired t test. Scale bars, 1 μm.

We further investigated whether *RPGRIP1L* mutations also affect the MKS complex components TMEM231 and TCTN2. TCTN2 and TMEM231 were both reduced within the transition zones of *RPGRIP1L* mutant cells (TCTN2 was reduced 56 ± 5%, n = [15–6, 15–4; 102, 80]; TMEM231 was reduced 62% ± 5%, n = [15–6, 15–4; 58, 50], Figure 5A). Thus, human RPGRIP1L is required for the transition zone localization of both NPHP1 and MKS complex components.

In addition to *RPGRIP1L* and *NPHP1*, mutations in *AHI1* cause JBTS^55,56^. We therefore examined whether RPGRIP1L affects the transition zone localization of AHI1. Like NPHP1, TCTN2, and TMEM231, established components of the NPHP and MKS complexes, AHI1 localized predominantly at the transition zone in a way that depended on RPGRIP1L (65 ± 6% reduction, n = [15–6, 15–4; 100, 69], Figure 5A).

To assess transition zone function, we examined whether JBTS-associated mutations in *RPGRIP1L* affect the ciliary localization of ARL13B. ARL13B levels were reduced by 49% ± 6%, n = (15–6, 15–4; 67, 33) in the fibroblasts of JBTS-affected individuals (Figure 5B). Thus, not only does JBTS-associated mutations in *TCTN2* (Figure 4A) disrupt the composition of the transition zone, JBTS-associated mutation of *RPGRIP1L* does as well, and both proteins are required to promote the ciliary localization of the JBTS-associated protein ARL13B.

Mutation of mouse *Rpgrip1l* leads to altered Hedgehog signaling^57^. As RPGRIP1L is required for the organization of the transition zone and the transition zone is a waypoint for SMO entering or leaving the cilium, we hypothesized that JBTS-associated mutations affect SMO localization to cilia. To test this hypothesis, we stimulated cells with the SMO agonist SAG and assessed SMO localization. In wild type human fibroblasts, SMO did not localize to cilia in the absence of SAG and accumulated within cilia in the presence of SAG, as expected (Figure 5C). In contrast, SAG stimulation only poorly induced the ciliary localization of SMO in fibroblasts of *RPGRIP1L* mutant cells; SMO levels in the cilium were reduced by 78% ± 9%, (n = [15–6 +SAG, 15–4 +SAG; 22, 21], Figure 5C). Moreover, we did not observe SMO docking within the transition zone in *RPGRIP1L* mutant fibroblasts (Figure 5C). Thus, JBTS-associated mutations in *RPGRIP1L* disrupt the composition of the transition zone, diminishing the ciliary localization of JBTS-associated proteins such as ARL13B and essential developmental regulators such as SMO.

### JBTS mutations in *RPGRIP1L* disrupt transition zone architecture

To further assess how RPGRIP1L functions in the human transition zone, we examined how JBTS-associated *RPGRIP1L* mutations affected the organization of the transition zone and ciliary membrane proteins. 3D STORM of control fibroblasts from an unaffected sibling and a JBTS-affected individual revealed that the JBTS-associated mutations in *RPGRIP1L* resulted in the loss of the RPGRIP1L ring in the transition zone (Figure 6A). To begin to examine the consequences of loss of transition zone RPGRIP1L, we assessed NPHP1 localization. We found that the NPHP1 ring was lost from the transition zones of *RPGRIP1L* mutant cells (Figure 6A). Thus, human RPGRIP1L is critical for the organization of the NPHP1 transition zone ring.

**Figure 6.**
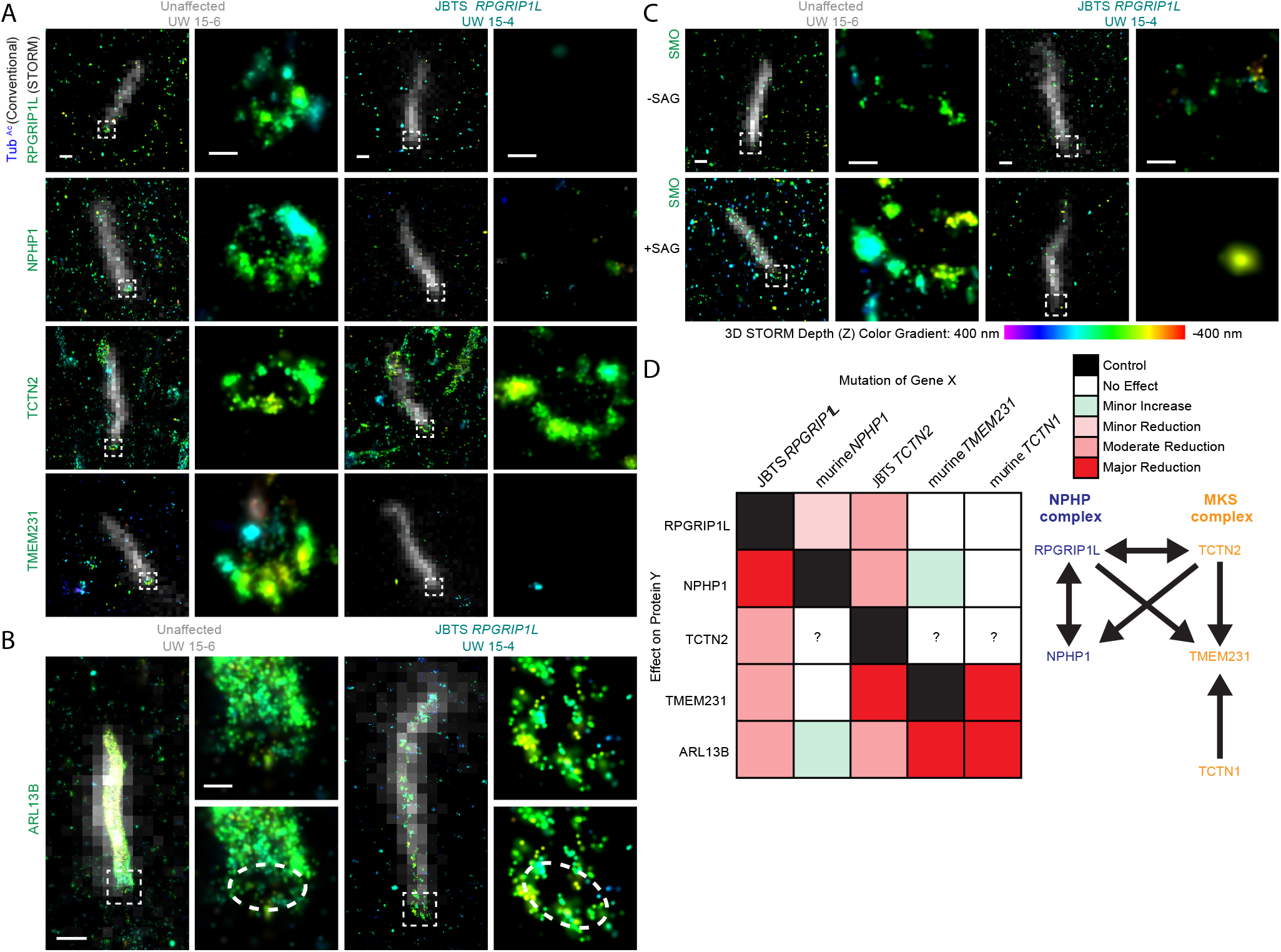
Transition zone rings formed by MKS and NPHP complexes are disrupted in a Joubert syndrome patient with mutations in *RPGRIP1L*. (**A**) Conventional epifluorescence images of acetylated tubulin (Tub^Ac^, white) overlayed on STORM images of RPGRIP1L, NPHP1, TCTN2, and TMEM231 (colored according to 3D STORM depth color gradient) in fibroblasts from a JBTS-affected individual with mutations in *RPGRIP1L* and an unaffected sibling. White boxes indicate location of the transition zone, magnified and displayed alongside the overlayed images. (**B**) Conventional epifluorescence images of acetylated tubulin (white) overlayed on STORM images of ARL13B (3D color gradient). Ring at the ciliary base is magnified and displayed alongside the overlayed images. Dashed line indicates ring pattern. (**C**) Conventional epifluorescence images of Tub^Ac^ (white) overlayed on STORM images of SMO in fibroblasts from a JBTS-affected individual and an unaffected sibling in the presence or absence of SAG. White boxes indicate location of the transition zone, magnified and displayed alongside the overlayed images. (**D**) Chart and model depicting the structural interdependence between the NPHP and MKS complexes. Left, mutation of *RPGRIP1L, NPHP1, TCTN2, TMEM231*^16,21^, *or TCTN1*^14^ has a specified effect on the levels of protein Y in the transition zone (RPGRIP1L, NPHP1, TCTN2, and TMEM231) or the cilium (ARL13B). ?, undetermined effect. Right, arrows indicating dependencies in transition zone protein localization. Scale bars, 500 nm for images overlayed with acetylated tubulin, and 100 nm for magnified images.

In addition to the NPHP ring, we investigated whether *RPGRIP1L* mutations affect the organization of the MKS ring components TMEM231 and TCTN2. STORM revealed that, although TCTN2 levels in the transition zone were reduced by ~60% in *RPGRIP1L* mutant cells (Figure 5A), the remaining TCTN2 formed a transition zone ring (Figure 6A). Thus, RPGRIP1L regulates the quantity of TCTN2 in the transition zone, but the organization of TCTN2 into a ring is independent of RPGRIP1L. In contrast, the TMEM231 ring was lost from the transition zones of *RPGRIP1L* mutant cells (Figure 6A). Thus, human RPGRIP1L is required for the transition zone localization of both NPHP1 and TMEM231, but TCTN2 can form into a ring independently of RPGRIP1L.

To assess transition zone function, we examined whether JBTS-associated mutations in *RPGRIP1L* affect the ciliary localization of ARL13B. In an unaffected individual, ARL13B was present throughout the ciliary membrane and as a ring in the transition zone (Figure 6B). In *RPGRIP1L* mutant cells, the transition zone ring of ARL13B was present but ARL13B was reduced in the cilium (Figure 6B). Thus, JBTS-associated mutation of *RPGRIP1L* only partially disrupts the function of the transition zone as a ciliary gate, or more specifically a ciliary waypoint.-Next we examined whether JBTS-associated mutations in *RPGRIP1L* affect the ciliary organization of SMO. In unaffected human fibroblasts, SMO did not localize to cilia in the absence of SAG and accumulated within cilia in the presence of SAG, as expected (Figure 6C). Furthermore, SMO docked within the transition zone forming a ring in the presence of SAG (Figure 6C). In contrast, SMO failed to accumulate in the cilium in the JBTS-affected individual, and SMO did not form a ring in the transition zone (Figure 6C). Thus, JBTS-associated mutations in *RPGRIP1L* disrupt the composition and architecture of the transition zone, diminishing the levels and impairing the organization of ciliary membrane proteins, such as ARL13B. We propose that some forms of JBTS are transition zone-opathies whose phenotypes are caused by compromised ciliary signaling secondary to disruption of the transition zone architecture.

## DISCUSSION

### Structural mapping by super-resolution microscopy reveals that JBTS-associated proteins form nested rings of doublets in the transition zone

Super-resolution microscopy has recently emerged as a powerful tool for analyzing the molecular organization of large protein complexes^28,31,58–60^. Key to this approach is determining the relative position of complex components. Subsequent calculation of positional coordinates can enable the assembly of a 3D rendering of a complex or, in our case, a sub-organellar compartment. Despite this capability, to date structural mapping by super-resolution microscopy has been mostly limited to one-dimensional position measurement linearly^58,59^ or radially^28,31,60^ organized structures. Here, we have demonstrated mapping of the ciliary base by analyzing the radial, angular and axial distribution of its components. To accomplish this mapping, we developed an angular distribution analysis method that is relatively insensitive to imperfect labeling efficiency and structural heterogeneities. We also demonstrated a self-registered, ratiometric two-color 3D STORM scheme that extracts channel registration information directly from the STORM raw data and determines the final position and color information from the two detection channels separately. Typically, two-color 3D STORM can be performed using either activator-reporter dye pairs^61^ or different reporter fluorophores^62^. The self-registration used here combines an advantage of the former -- avoidance of the need for laborious 3D channel registration -- with an advantage of the latter -- minimal channel crosstalk. Both the imaging and analysis methods can be readily applied to other structural mapping efforts.

By determining the arrangement of key ciliopathy-associated proteins in the transition zone and the distal appendages using super-resolution microscopies, we found that the MKS and NPHP complexes form nested rings, with the MKS complex positioned at the transition zone membrane and the NPHP complex positioned between the ciliary axoneme and the ciliary membrane. Both of the NPHP and MKS complexes form nine-fold doublets, closely matching the doublets formed by the arms of the electron micrographically defined Y links. We found that human mutations in *RPGRIP1L* and *TCTN2* that cause JBTS decrease both the NPHP and MKS complex at the transition zone. In addition, identifying that proteins of the NPHP and MKS complexes exhibit a similar doublet arrangement and are in close proximity suggests that RPGRIP1L and TCTN2, and by extension the NPHP and MKS complexes, have interdependent functions.

The shared arrangement of the MKS and NPHP complexes and their striking similarity to the position and conformation of the Y links suggests that they are components of the arms of the Y links. Another transition zone component, CEP290, underlies JBTS, MKS, and NPHP^63–65^ and, in *Chlamydomonas*, its homolog localizes to the Y links^26^. The co-localization of multiple protein complexes within the transition zone is paralleled by their shared functions in promoting ciliogenesis and controlling ciliary composition^14,16,21^. Our finding that JBTS-causing mutations in *RPGRIP1L* and *TCTN2* disrupt both the NPHP and MKS complexes in the transition zone reveals the structural basis for the functional interactions that have been observed between the two complexes. For example, in mice, compound mutation of the MKS complex gene *Tctn1* and either *Nphp1* or *Nphp4* heightens defects in ciliogenesis and Hedgehog developmental signaling compared to the individual mutations^66^. Similarly, in *C. elegans*, mutation of individual MKS or NPHP complex genes do not disrupt ciliogenesis, but compound mutation of MKS and NPHP complex genes abrogate ciliary structure^23,66^. The overlapping localizations and functions of the NPHP and MKS complexes suggest that they have partially overlapping roles in controlling ciliogenesis and ciliary composition (Figure 6D). Our structural analysis suggests that this functional overlap may reflect partially interdependent generation of the transition superassembly by the MKS and NPHP complexes.

### The transition zone is a waypoint for diverse ciliary proteins

Mutation of any of several transition zone MKS complex components disrupts the ciliary localization of structurally diverse ciliary membrane-associated proteins^14,16,21^. Disruption of the transition zone also allows normally non-ciliary membrane proteins to leak into the cilium^21–23^. Thus, the transition zone acts as a gate to allow some proteins to accumulate within the cilium, either by facilitating ciliary entry or preventing ciliary exit, and restrict the entrance of others. This gating is essential for ciliary signaling and vertebrate development. For example, Hedgehog binding to PTCH1 allows SMO to accumulate at the cilium to activate the downstream Gli transcription factors^67^. We found that upon activation of the Hedgehog pathway, SMO accumulates at discrete clusters in the transition zone. Single-molecule imaging has shown that SMO exhibits primarily diffusive movement in the cilium with binding events in the vicinity of the ciliary base^36^. Using two-color STORM, we observed that these binding sites at the ciliary base are in the transition zone, highlighting the importance of the transition zone in regulating SMO trafficking to the cilium. As SMO localization to the transition zone requires RPGRIP1L, RPGRIP1L or RPGRIP1L-dependent proteins likely form the SMO binding sites. This accumulation in the transition zone, together with the requirement for the MKS complex for ciliary localization, suggests that the transition zone acts as a way station for SMO on its entry or exit into or out of the cilium.

Previous studies have suggested that the transition zone may also be a diffusion barrier to prevent the ciliary exit of proteins like SMO. However, the distances between the transition zone complexes analyzed here suggests that proteins smaller than 50 nm could diffuse between them. As SMO has a width of 5 nm^68^, if the transition zone functions as a diffusion barrier, it must rely on proteins beyond those characterized here.

Why might ciliary proteins pause at the transition zone? One possibility is that the transition zone is the place where ciliary cargo is loaded onto the intraflagellar transport machinery entering the cilium. Consistent with the model of coordinated trafficking through the transition zone, IFT88 also accumulated at the transition zone (Figures 3H and I).

### Joubert syndrome can be caused by transition zone disruption

The involvement of ciliary proteins in both Hh signaling and JBTS have suggested that disruptions in Hh signaling may underlie JBTS^69,70^. Our data reveal that mutations in human *TCTN2* that cause JBTS disrupt the localization of other JBTS-associated transition zone proteins including NPHP1, RPGRIP1L, and TMEM231. Moreover, mutations in human *RPGRIP1L* that cause JBTS disrupt the localization of JBTS-associated transition zone proteins NPHP1, TCTN2, TMEM231 and AHI1. Still other JBTS-associated transition zone proteins (*e.g*., CEP290, TMEM67, CC2D2A, TCTN1, TCTN3, B9D1 and MKS1) may well also depend on TCTN2 and RPGRIP1L for localization. Of interest, the requirement for human RPGRIP1L in localizing both NPHP and MKS components parallels the requirement for a RPGRIP1L homolog, MKS-5, in *C. elegans*^22,23^. Thus, RPGRIP1L is a core organizer of the transition zone whose functions are evolutionarily conserved from nematodes to humans. In contrast, the sole member of the *C. elegans* Tectonic family is dispensable for MKS-5 transition zone localization^66^, indicating that the functions of other transition zone components may have changed over evolution.

We have previously reported that null mutations in genes encoding MKS complex components cause defects in murine ciliary membrane composition, including SMO, and are associated with disrupted Hh signaling^14,20^. Here, we examined the effects of compound heterozygous JBTS-associated mutations in human *RPGRIP1L* and found that, like the MKS complex, RPGRIP1L is essential for SMO accumulation at the cilium in response to pathway activation. It seems likely that RPGRIP1L, by localizing the MKS complex to the transition zone, allows SMO to accumulate at the cilium and activate the downstream Hh pathway.

Unlike most JBTS-associated proteins, ARL13B localizes to the ciliary membrane. We found that JBTS-associated mutations in *TCTN2* and *RPGRIP1L* compromise localization of ARL13B to the cilium. Thus, the diverse genes underlying JBTS may fall into one of at least three categories: 1) genes, such as *TCTN2* and *RPGRIP1L*, that encode core components required for organizing the transition zone, 2) genes, such as *NPHP1*, that encode components of the transition zone not essential for its organization but contributing to its function, and 3) genes, such as *ARL13B*, that require the transition zone for ciliary localization. The generation of the transition zone super-assembly is critical for the localization of select membrane-associated proteins such as ARL13B and SMO to cilia, and perturbation of its architecture is one of the causes of JBTS.

## ACKNOWLEDGEMENTS

Structured illumination microscopy was performed on a Nikon N-SIM system in the UCSF Nikon Imaging Center. We thank Harrison Liu and Joerg Schnitzbauer for help setting up the STORM system and Joerg Schnitzbauer in developing the analysis algorithms. We thank Vicente Herranz-Pérez and Jose Manuel Garcia-Verdugo for providing electron micrographs. This project is supported by the NIH Director’s New Innovator Award (DP2OD008479) to X.S., R.M. and B.H. and by grants from the NIH (AR054396 and GM095941 to J.F.R, F32GM109714 to G.G., and U54HD083091 sub-project 6849 to D.D.) and the Burroughs Wellcome Fund and the Packard Foundation to G.G. and J.F.R. B.H. is a Chan Zuckerberg Biohub investigator.

## AUTHOR CONTRIBUTIONS

X.S., G.G., B.H. and J.F.R. designed the experiments and wrote the manuscript, X.S. and R.M. built the STORM microscope, G.G. generated the samples, X.S. and G.G. performed the STORM imaging experiments, and X.S. analyzed the STORM data. G.G. performed the SIM imaging experiments and analyzed the SIM data. J.C.W. and D.D. collected and genotyped the human fibroblasts and provided feedback on the manuscript. G.J.P. generated the α-NPHP1 antibody.

## METHODS

### Primary mouse tracheal epithelial cell culture

Primary mouse tracheal epithelial cultures were derived as described^71^. In summary, tracheas from C57BL/6J mice aged 8 to 12 weeks were dissected. The tracheal cells were dissociated with protease. Dissociated cells were placed in a culture plate (Primaria) for several hours to remove fibroblasts. The non-adherent tracheal epithelial cells were seeded at a density of 0.33 × 10^5^ cells per 24-well insert (Day 0). The cells were cultured for three days with mTEC/Plus medium in the basal and apical compartments of the transwell. After three days of incubation (Day 3), the medium was replaced with fresh mTEC/Plus medium. Air-liquid interface (ALI) differentiation was begun at day 5 by removing the medium in the apical transwell chamber and replacing the medium in the basal chamber with mTEC/NS. The medium was replaced every other day with mTEC/NS, and the filters were harvested at ALI day 14 for immunofluorescence experiments. All animal protocols were approved by the Institutional Animal Care and Use Committee (IACUC) of the University of California, San Francisco.

### IMCD3 and fibroblast cell culture

IMCD3 cells were cultured in DMEM:F12 supplemented with 10% FBS. MEFs and human primary dermal fibroblasts were cultured in DMEM, Glutamax (Thermo Fisher), and 10% FBS. For imaging, IMCD3 cells, MEFs, and human primary dermal fibroblasts were seeded at 2.5 × 10^4^ cells/cm^2^ and grown to confluency. Cells were starved of serum in Opti-MEM reduced serum medium for 24 hours to induce ciliogenesis.

### Fixation and staining for immunofluorescence

mTEC filters were harvested at ALI day 14 and the apical and basal compartments were washed with phosphate buffered saline (PBS) three times. The filters were fixed by adding 4% paraformaldehyde (PFA) in PBS to the apical compartment and incubating at room temperature for ten minutes. The PFA was removed by washing the filter with PBS six times with at least one minute of incubation between washes. The filters were then treated with a solution of 1% sodium dodecyl sulfate (SDS) for five minutes in PBS to reveal antigens. The SDS solution was removed by washing the filter with PBS.

For SIM, filters were incubated in blocking buffer (PBS, 2.5 % BSA, 0.1 % Triton X-100, 0.02 % sodium azide) for ten minutes at room temperature. Primary antibodies were added to the filters in blocking buffer according as described in Table 1. The filters were incubated overnight (~16 hours) at 4 degrees on a rocker. The filters were washed with PBT (PBS, 0.2% Triton X-100) six times with at least one minute of incubation between washes. Secondary antibodies were added at a dilution of 1:1000 and incubated for two hours in blocking buffer on an orbital shaker. The secondary antibody was removed by six washes with PBT buffer. The filters were mounted on slides in Prolong Gold containing 4',6-diamidino-2-phenylindole (DAPI). High performance coverslips were used, and the mounting medium was allowed to solidify for 16–24 hours before sealing the slides with nail polish and imaging.

For STORM, filters were incubated in blocking buffer for ten minutes at room temperature. Primary antibodies were added to the filters incubating in blocking buffer according to the table of antibodies and concentrations (Table 1). The filters were incubated with the primary antibody overnight (~16 hours) at 4 °C on a rocker. The filter was washed with PBT. Secondary antibodies were incubated for two hours at room temperature or overnight (~16 hours) at 4 °C on the filters in blocking buffer. The secondary antibody was removed by six washes with PBT.

Some samples for single color STORM were stained using labeled primary antibodies. In these cases, filters were incubated in blocking buffer for ten minutes at room temperature. Primary antibodies were added to the filters in blocking buffer at concentrations listed in Table 1. The filters were incubated with the primary antibody overnight (~16 hours) at 4 °C on a rocker. The filter was washed with PBT.

For two-color STORM using primaries derived from the same species, filters were incubated in IF blocking buffer for ten minutes at room temperature. Primary antibody recognizing the experimental protein was added to the filters incubating in blocking buffer. The filters were incubated with the primary antibody overnight (~16 hours) at 4 °C on a laboratory rocker. The filter was washed with PBT. Secondary antibody labelled with Alexa 647 was added according to the table and incubated for two hours at room temperature on the filters in IF blocking buffer. The secondary antibody was removed by six washes with Wash buffer. Primary antibody labelled with Cy5.5 recognizing the reference protein was added to the filters incubating in blocking buffer according to the table of antibodies and concentrations. The filters were incubated with the primary antibody overnight (~16 hours) at 4 °C on a laboratory rocker. The filter was washed with PBT.

### Mounting filters in STORM buffer

After immunofluorescence staining, mTEC filters were mounted on slides and high-performance coverslips in PBS with the addition of 100 mM mercaptoethylamine at pH 8.5, 5% glucose (wt/vol) and oxygen scavenging enzymes [0.5 mg/ml glucose oxidase (Sigma-Aldrich), and 40 mg/ml catalase (Roche Applied Science)]. The buffer remains suitable for imaging for two hours.

### SIM image acquisition

SIM data were collected on a N-SIM microscope in 3D-SIM mode using an Apo TIRF 100x oil NA 1.49 objective (Nikon) following standard operation procedures. Image reconstruction was using the NIS Elements software (Nikon).

### STORM instrumentation

STORM experiments were performed on a custom-built microscope based on a Nikon Ti-U inverted microscope. Three activation/imaging photodiode lasers (Coherent CUBE 405, OBIS 561 and CUBE 642) were combined using dichroic mirrors, aligned, expanded and focused to the back focal plane of the objective (Nikon Plan Apo 100× oil NA 1.45). The lasers were controlled directly by the computer. A quadband dichroic mirror (zt405/488/561/640rpc, Chroma) and a band-pass filter (ET705/70m, Chroma) separated the fluorescence emission from the excitation light. The focus drift of the microscope sample stage during image acquisition was stabilized by a closed-loop focus stabilization system that sends an infra-red laser beam through the edge of the microscope objective and detects the back reflection from the sample coverglass with a CCD camera^35^.

After the emission light exits the microscope side port, we added a low-end piezoelectric deformable mirror (DM) (DMP40-P01, Thorlabs) at the conjugate plane of the objective pupil plane. This DM contains 40 actuators arranged in a circular keystone pattern. By first flattening the mirror and then manually adjusting key Zernike polynomials of the DM surface, we found that this DM has sufficient performance to correct aberrations induced by the optical system. We did not observe obvious depth-induced spherical aberrations caused by glass-water refractive index mismatch because transition zones we imaged were < 1 μm away from the coverglass surface. Moreover, after correcting the intrinsic aberrations, we increased the primary astigmatism Zernike coefficient of the DM surface to introduce astigmatism for 3D STORM. To experimentally generate a calibration curve of point spread function as a function of z, we immobilized Alexa 647–labeled antibodies on a coverglass and imaged individual molecules as the sample was scanned in z^35^. Compared to using a cylindrical lens^35^ using the DM to introduce astigmatism did not cause magnification differences in the two lateral directions. It also allowed tractable optimization of the amount of astigmatism to achieve the best axial resolution. The Z localization precision (Z resolution) is estimated to be ~70 nm full width at half maximum (FWHM) based on the reconstructed STORM images. The Z focal depth is ~800 nm. We measured the longitudinal distance between the transition zone and distal appendages by analyzing only transition zone structures with their axes close to the XY plane. Therefore, the precision was limited mostly by the XY resolution instead of Z resolution.

The fluorescence was subsequently split by a 705 nm beam splitter (T705lpxr, Chroma) to a long wavelength channel (ch1, >705 nm) and a short wavelength channel (ch2, <705 nm). The two channels were recorded at a frame rate of 57 Hz on an electron multiplying CCD camera (Ixon+ DU897E-CS0-BV, Andor). The split-view setup discriminates fluorescent labeled by small shifts (~20 nm) in their emission spectra, in this work Alexa Fluor 647 and Cy5.5. Because individual emitters were recorded in both long and short channels, the intensity ratio between the channels is sufficient to identify each emitter^72^.

### STORM image acquisition and analysis

Photoswitchable dye pairs Alexa 405/Alexa 647 and Cy3/Cy5.5 were used for STORM imaging with a ratio of 0.8 Alexa 647 molecules per antibody, and 0.5 Cy5.5 per antibody. During the imaging process, the 405 nm and 561 nm activation lasers were used to activate a small fraction of the Alexa 647 and Cy5.5 reporters at a time, and individual activated fluorophores were excited with a 642 nm laser. The typical power for the lasers at the back port of the microscope was 30 mW for the 642 nm imaging laser and 0.5 to 5 μW for activation lasers.

Analysis of STORM raw data was performed in the Insight3 software^35^, which identifies and fits single molecule spots in each camera frame to determine their x, y and z coordinates as well as photon numbers. Sample drift during data acquisition were corrected using imaging correlation analysis. The drift-corrected coordinates, photon number, and the frame of appearance of each identified molecule were saved in a molecule list for further analysis.

To estimate position uncertainty, we considered several factors. First, we measured singlemolecule localization precision in our images to be ~15 nm (SD) by analyzing the relative position of the same photoactivated molecule appearing in consecutive frames. Second, the antibody size contributed additional error. Previous STORM of indirectly immunostained microtubules revealed that primary and secondary antibodies adds ~30 nm to the overall diameter^34^, suggesting a localization bias up to 15 nm in cases in which primary and secondary antibodies bind from a uniform direction. Given that we used polyclonal antibodies of all TZ targets, we did not anticipate biased binding orientation to occur. When the orientation of antibody binding is random, combining the 15 nm localization precision and a 30 nm uniform distribution from the antibodies (SD = 9 nm) results in an overall uncertainty of 17 nm (SD).

### Ratiometric color identification and self-referenced registration

Because of the high labeling density for transition zone proteins, multi-color STORM utilizing activator-reporter dye pairs with different activation wavelengths^34^ suffer from high cross-talk between color channels. Therefore, we choose to use reporter dyes with distinct emission spectra for two-color imaging. Typically, this approach uses a dichroic mirror to split the emission light into two distinct channels, most commonly two halves of the same camera chip. The drawback, though, is that the two channels must be registered with nanometer accuracy. This registration requires additional calibration images acquired before every imaging experiment (e.g. using fluorescent beads). Moreover, three-dimensional channel registration could be complicated due to the chromatic aberrations between channels.

To overcome these inconveniences, we purposely used two reporter dyes which have substantial overlap in the emission spectra. We chose far-red photoswitchable cyanine dyes Alexa Fluo 647 and Cy5.5 because they have superior photoswitching characteristics than most shorter-wavelength dyes. In our system, approximately 60% of Alexa 647 emission and 48% of Cy5.5 emission went into the shorter wavelength channel and the rest into the longer wavelength channel. Because the same Cy5.5 molecule appears in both channels with approximately equal intensity, the STORM data itself contains channel registration information. We extracted this information by first manually identifying 3–4 pairs matched spots in the two channels to calculate crude registration information, then automatically identify all matched spots in ~ 1000 STORM frames using this crude registration information, and finally determine a polynomial coordinate transform function between the two channels with a least square fitting to the coordinates of these matched spots. The least square fitting can be performed for just the xy coordinates or for all xyz coordinates. In this work, we only performed 2D registration for the subsequent determination of color identity. The polynomial transform function is: x_c_ = A_0_ + A_1_ x + B_1_ y + A_2_ x^2^ + C_2_ x y + B_2_ y^2^ + A_3_ x^3^ + C_31_ x y^2^ + C_32_ x^2^ y + B_3_ y^3^; y_c_ = E_0_ + E_1_ y + F_1_ x + E_2_ y^2^ + G_2_ x y + F_2_ x^2^ + E_3_ y^3^ + G_32_ x^2^ y + G_31_ x y^2^ + F_3_ x^3^; where x_c_ and y_c_ are the transformed coordinates, x and y are the initial coordinates, and A_0_, A_1_, A_2_, A_3_, B_1_, B_2_, B_3_, C_2_, C_31_, C _32_, E_0_, E_1_, E_2_, E_3_, F_1_, F_2_, F_3_, G_2_, G_31_, and G_32_ are coefficients obtained by the least square fitting.

With the 2D registration information, all spots identified in the short wavelength channel (containing both Alexa 647 and Cy5.5) were mapped to the longer wavelength cannel to calculate the corresponding intensity. The color identity of each single molecule spot was then determined based on the intensities in the two channels, *I*_S_ and *I*_L_. Using samples labeled with only one of the two dyes, we generated a calibration plot in the log(*I*_S_)−log(*I*_L_) representation, with areas corresponding to the two dyes cleanly separated by a straight line. The crosstalk between the two channels is approximately 15%.

To construct the final super-resolution image, we used the coordinate information from only the shorter wavelength channel so that the two colors are perfectly aligned without chromatic aberration. Although this approach sacrifices spatial resolution because some of the emission photons are in the longer wavelength detection channel, we found that the final spatial resolution was sufficient for our study. In applications demanding better spatial resolution, it would be straightforward to merge the positional information from both channels utilizing the channel registration information extracted from the STORM data itself.

### Diameter measurement

We robust-fitted each ring structure imaged by STORM to an ellipse^37^. We chose robust fit instead of least square fit to avoid outlier effects from noise localization points in the STORM image. The diameter of a circle with the same perimeter of the ellipse was considered as the diameter of its corresponding structure. The diameter of structure formed by each kind of protein was determined by averaging about 30 fitted structures. The proteins we measured were NPHP1, TEM231, B9D1 and RPGRIP1L in the transition zone, and CEP164 in distal appendages.

### Image alignment using discrete Fourier transform (DFT) and cross correlation

The alignment algorithm is a modified version of image registration^73^ that achieved efficient subpixel image registration by upsampled DFT cross correlation and takes into account sample rotation. Briefly, the original algorithm (1) obtains an initial 2D shift estimate of the cross correlation peak by fast Fourier transforming (FFT) a image to register and a reference image, (2) and then refines the shift estimation by upsampling the DFT only in a small neighborhood of that estimate by means of a matrix-multiply DFT. We modified step (1) by rotating the image to align from 0 to 359° by 1° at a step, and then performing original step (1) after each rotation step to obtain 360 cross correlation peaks. The biggest peak provides not only the initial 2D shift estimate but also the optimal angle of rotation. The initial 2D shift estimate then goes through the same step (2) in the original algorism to achieve subpixel alignment. The images of single structure were aligned for 10 iterations. The superimposition of all images was used as the reference image for the first iteration. Each image was translated and rotated to the reference image to maximize the cross correlation between the image to align and reference image in Fourier domain. The superimposition of the images aligned in one iteration was used as the reference image for the next iteration. Our algorithm aligned the images within 1/100 of a pixel.

### Nearest-cluster angular spacing analysis and bootstrapping

The nearest-cluster angular spacing analysis is based on the coordinates in STORM images. To analyze the angles between nearest clusters in the transition zone protein and CEP164 rings, we (1) fitted each structure to a ring, converted the Cartesian coordinates to polar coordinates, and constructed a histogram of the angles of all localizations in each STORM image, (2) found the peaks in the histogram with multiple Gaussian functions using a matlab program findpeaksb.m developed by Dr. T. C. O'Haver at Univeristy of Maryland (https://terpconnect.umd.edu/~toh/spectrum/PeakFindingandMeasurement.htm), (3) calculated the angle between nearest peaks, (4) plotted a histogram of the central angles between nearest peaks (clusters) obtained from n individual images (n = 50~125), and finally (5) found the peaks in the histogram of nearest-cluster angular spacing by fitting them to multiple Gaussian functions using findpeaksb.m (Figure 2E-H). The positions of the centers of the fitted Gaussian functions obtained in step (5) were used as the angles between adjacent clusters in the model of TZ architecture (Figure 2E-H). The uncertainties of these angles were estimated by bootstrapping analysis. This technique allows estimation of the sampling distribution by random sampling the individual images to new sets of images. Each new set has the same number of images as in the original set. We randomly sampled 200 sets of images for each protein in our bootstrapping analysis to test the robustness of peak identification and calculate the standard deviation of nearest-cluster angles. For TZ proteins NPHP1, TMEM231, B9D1, and RPGRIP1L, more than 80% of the bootstrapping histograms of nearest-cluster angles have three peaks as demonstrated in Figures 2E-H.

The findpeaksb.m program determines the number of peaks and the position, width and height of each peak by least-squares curve-fitting to the original histograms. More specifically, it smooths the first derivative of the histogram, detects downward-going zero-crossings, and takes only those zero crossings whose slope exceeds a certain predetermined minimum (called the slope threshold) t a point where the original signal exceeds a certain minimum (called the amplitude threshold"). We set the smooth width, slope threshold, and amplitude threshold for the algorithm, and kept these parameters constant for CEP164 (Figure 2E) and transition zone proteins (Figure 2F-H).

### Quantitation of fluorescence intensity in human fibroblasts

Confocal images of cilia were collected on a Leica SPE laser-scanning confocal equipped with four laser lines (405 nm, 488 nm, 551 nm, and 635 nm) and an ACS APO 63x/1.30 NA oil objective lens. The fluorescence intensity of individual cilia and transition zones in confocal micrographs was measured using Fiji software^74^. The average background fluorescence intensity in the vicinity of each transition zone and cilium was subtracted from the fluorescence intensity of individual transition zones and cilia.

### Structural homology modelling

Homology models were generated using ModWeb version r182^75,76^.

### Domain architecture prediction

Domain architecture predications were generated using JPred 4 and Topcons web servers^77,78^.

### Statistics and reproducibility

Statistical analyses were performed using Graphpad Prism 7.0a. Graphical data are presented as mean ± 95% confidence interval or mean ± standard deviation as specified in the figure legends. n values are stated in the figure legends and definition of n is number of transition zones (Figures 4A, 4C, and 5A) or number of cilia (Figures 4B, 4D, 5B, and 5C). P values are stated in the figure legends, and symbols indicate the following P values: ns, P > 0.05. *, P ≤ 0.05. **, P ≤ 0.01. ***, P ≤ 0.001. ****, P ≤ 0.0001. Two sample comparisons were performed using an unpaired t test with a two tailed P value. Multiple comparisons were performed using an ordinary one-way ANOVA, along with Tukey’s multiple comparisons test to analyze pairwise differences. Fluorescence intensity measurements (Figures 4 and 5) from multiple experiments were combined for statistical analysis. Negative normalized fluorescence intensity values were set to zero in graphs. Experiments were independently performed either twice (Figures 4A, 4B, and 5C) or thrice (Figure 4C, 5A, and 5B).

## REFERENCES

1. Dutcher, S. K. The awesome power of dikaryons for studying flagella and basal bodies in Chlamydomonas reinhardtii. Cytoskeleton (Hoboken) 71, 79–94 (2014).

2. Pearson, C. G. & Winey, M. Basal body assembly in ciliates: the power of numbers. Traffic 10, 461–471 (2009).

3. Garcia, G. & Reiter, J. F. A primer on the mouse basal body. Cilia 5, 17 (2016).

4. Vertii, A., Hung, H.-F., Hehnly, H. & Doxsey, S. Human basal body basics. Cilia 5, 13 (2016).

5. Anderson, R. G. The three-dimensional structure of the basal body from the rhesus monkey oviduct. J Cell Biol 54, 246–265 (1972).

6. Gilula, N. B. & Satir, P. The ciliary necklace. A ciliary membrane specialization. J Cell Biol 53, 494–509 (1972).

7. Habbig, S. & Liebau, M. C. Ciliopathies - from rare inherited cystic kidney diseases to basic cellular function. Mol Cell Pediatr 2, 8 (2015).

8. Szymanska, K., Hartill, V. L. & Johnson, C. A. Unraveling the genetics of Joubert and Meckel-Gruber syndromes. J Pediatr Genet 3, 65–78 (2014).

9. Otto, E. et al. A gene mutated in nephronophthisis and retinitis pigmentosa encodes a novel protein, nephroretinin, conserved in evolution. Am J Hum Genet 71, 1161–1167 (2002).

10. Wolf, M. T. F. et al. Mutational analysis of the RPGRIP1L gene in patients with Joubert syndrome and nephronophthisis. Kidney Int. 72, 1520–1526 (2007).

11. Srour, M. et al. Mutations in TMEM231 cause Joubert syndrome in French Canadians. J Med Genet 49, 636–641 (2012).

12. Arts, H. H. et al. Mutations in the gene encoding the basal body protein RPGRIP1L, a nephrocystin-4 interactor, cause Joubert syndrome. Nat Genet 39, 882–888 (2007).

13. Hopp, K. et al. B9D1 is revealed as a novel Meckel syndrome (MKS) gene by targeted exon-enriched next-generation sequencing and deletion analysis. Hum Mol Genet 20, 2524–2534 (2011).

14. Garcia-Gonzalo, F. R. et al. A transition zone complex regulates mammalian ciliogenesis and ciliary membrane composition. Nat Genet 43, 776–784 (2011).

15. Dowdle, W. E. et al. Disruption of a ciliary B9 protein complex causes Meckel syndrome. Am J Hum Genet 89, 94–110 (2011).

16. Roberson, E. C. et al. TMEM231, mutated in orofaciodigital and Meckel syndromes, organizes the ciliary transition zone. J Cell Biol 209, 129–142 (2015).

17. Delous, M. et al. The ciliary gene RPGRIP1L is mutated in cerebello-oculo-renal syndrome (Joubert syndrome type B) and Meckel syndrome. Nat Genet 39, 875–881 (2007).

18. Joubert, M., Eisenring, J. J., Robb, J. P. & Andermann, F. Familial agenesis of the cerebellar vermis. A syndrome of episodic hyperpnea, abnormal eye movements, ataxia, and retardation. Neurology 19, 813–825 (1969).

19. Parisi, M. A., Doherty, D., Chance, P. F. & Glass, I. A. Joubert syndrome (and related disorders) (OMIM 213300). Eur. J. Hum. Genet. 15, 511–521 (2007).

20. Sang, L. et al. Mapping the NPHP-JBTS-MKS protein network reveals ciliopathy disease genes and pathways. Cell 145, 513–528 (2011).

21. Chih, B. et al. A ciliopathy complex at the transition zone protects the cilia as a privileged membrane domain. Nat Cell Biol 14, 61–72 (2012).

22. Jensen, V. L. et al. Formation of the transition zone by Mks5/Rpgrip1L establishes a ciliary zone of exclusion (CIZE) that compartmentalises ciliary signalling proteins and controls PIP2 ciliary abundance. EMBO J 34, 2537–2556 (2015).

23. Williams, C. L. et al. MKS and NPHP modules cooperate to establish basal body/transition zone membrane associations and ciliary gate function during ciliogenesis. J Cell Biol 192, 1023–1041 (2011).

24. Reiter, J. F., Blacque, O. E. & Leroux, M. R. The base of the cilium: roles for transition fibres and the transition zone in ciliary formation, maintenance and compartmentalization. EMBO Rep 13, 608–618 (2012).

25. Awata, J. et al. NPHP4 controls ciliary trafficking of membrane proteins and large soluble proteins at the transition zone. J Cell Sci 127, 4714–4727 (2014).

26. Craige, B. et al. CEP290 tethers flagellar transition zone microtubules to the membrane and regulates flagellar protein content. J Cell Biol 190, 927–940 (2010).

27. Garcia-Gonzalo, F. R. & Reiter, J. F. Scoring a backstage pass: mechanisms of ciliogenesis and ciliary access. J Cell Biol 197, 697–709 (2012).

28. Mennella, V. et al. Subdiffraction-resolution fluorescence microscopy reveals a domain of the centrosome critical for pericentriolar material organization. Nat Cell Biol 14, 1159–1168 (2012).

29. Lau, L., Lee, Y. L., Sahl, S. J., Stearns, T. & Moerner, W. E. STED microscopy with optimized labeling density reveals 9-fold arrangement of a centriole protein. Biophys J 102, 2926–2935 (2012).

30. Yang, T. T. et al. Superresolution Pattern Recognition Reveals the Architectural Map of the Ciliary Transition Zone. Sci Rep 5, 14096 (2015).

31. Lawo, S., Hasegan, M., Gupta, G. D. & Pelletier, L. Subdiffraction imaging of centrosomes reveals higher-order organizational features of pericentriolar material. Nat Cell Biol 14, 1148–1158 (2012).

32. Gustafsson, M. G. Surpassing the lateral resolution limit by a factor of two using structured illumination microscopy. J Microsc 198, 82–87 (2000).

33. Rust, M. J., Bates, M. & Zhuang, X. Sub-diffraction-limit imaging by stochastic optical reconstruction microscopy (STORM). Nat Methods 3, 793–795 (2006).

34. Bates, M., Huang, B., Dempsey, G. T. & Zhuang, X. Multicolor super-resolution imaging with photo-switchable fluorescent probes. Science 317, 1749–1753 (2007).

35. Huang, B., Wang, W., Bates, M. & Zhuang, X. Three-dimensional super-resolution imaging by stochastic optical reconstruction microscopy. Science 319, 810–813 (2008).

36. Milenkovic, L. et al. Single-molecule imaging of Hedgehog pathway protein Smoothened in primary cilia reveals binding events regulated by Patched1. Proc Natl Acad Sci USA 112, 8320–8325 (2015).

37. Taubin, G. Estimation of Planar Curves, Surfaces, and Nonplanar Space-Curves Defined by Implicit Equations with Applications to Edge and Range Image Segmentation. IEEE Transactions on Pattern Analysis and Machine Intelligence 13, 1115–1138 (1991).

38. Alcedo, J., Ayzenzon, M., Ohlen Von, T., Noll, M. & Hooper, J. E. The Drosophila smoothened gene encodes a seven-pass membrane protein, a putative receptor for the hedgehog signal. Cell 86, 221–232 (1996).

39. Rohatgi, R., Milenkovic, L. & Scott, M. P. Patched1 regulates hedgehog signaling at the primary cilium. Science 317, 372–376 (2007).

40. Corbit, K. C. et al. Vertebrate Smoothened functions at the primary cilium. Nature 437, 1018–1021 (2005).

41. Hahn, H. et al. Mutations of the human homolog of Drosophila patched in the nevoid basal cell carcinoma syndrome. Cell 85, 841–851 (1996).

42. Johnson, R. L. et al. Human homolog of patched, a candidate gene for the basal cell nevus syndrome. Science 272, 1668–1671 (1996).

43. Goodrich, L. V., Milenkovic, L., Higgins, K. M. & Scott, M. P. Altered neural cell fates and medulloblastoma in mouse patched mutants. Science 277, 1109–1113 (1997).

44. Raffel, C. et al. Sporadic medulloblastomas contain PTCH mutations. Cancer Res 57, 842–845 (1997).

45. Chen, J. K., Taipale, J., Young, K. E., Maiti, T. & Beachy, P. A. Small molecule modulation of Smoothened activity. Proc Natl Acad Sci USA 99, 14071–14076 (2002).

46. Ou, Y. et al. Adenylate cyclase regulates elongation of mammalian primary cilia. Exp Cell Res 315, 2802–2817 (2009).

47. Bishop, G. A., Berbari, N. F., Lewis, J. & Mykytyn, K. Type III adenylyl cyclase localizes to primary cilia throughout the adult mouse brain. J Comp Neurol 505, 562–571 (2007).

48. Wong, S. T. et al. Disruption of the type III adenylyl cyclase gene leads to peripheral and behavioral anosmia in transgenic mice. Neuron 27, 487–497 (2000).

49. Livera, G. et al. Inactivation of the mouse adenylyl cyclase 3 gene disrupts male fertility and spermatozoon function. Mol. Endocrinol. 19, 1277–1290 (2005).

50. Wang, Z. et al. Adult type 3 adenylyl cyclase-deficient mice are obese. PLoS ONE 4, e6979 (2009).

51. Deane, J. A., Cole, D. G., Seeley, E. S., Diener, D. R. & Rosenbaum, J. L. Localization of intraflagellar transport protein IFT52 identifies basal body transitional fibers as the docking site for IFT particles. Curr Biol 11, 1586–1590 (2001).

52. Stratigopoulos, G. et al. Hypomorphism for RPGRIP1L, a ciliary gene vicinal to the FTO locus, causes increased adiposity in mice. Cell Metab. 19, 767–779 (2014).

53. Parisi, M. A. et al. The NPHP1 gene deletion associated with juvenile nephronophthisis is present in a subset of individuals with Joubert syndrome. Am J Hum Genet 75, 82–91 (2004).

54. Cantagrel, V. et al. Mutations in the cilia gene ARL13B lead to the classical form of Joubert syndrome. Am J Hum Genet 83, 170–179 (2008).

55. Dixon-Salazar, T. et al. Mutations in the AHI1 gene, encoding jouberin, cause Joubert syndrome with cortical polymicrogyria. Am J Hum Genet 75, 979–987 (2004).

56. Ferland, R. J. et al. Abnormal cerebellar development and axonal decussation due to mutations in AHI1 in Joubert syndrome. Nat Genet 36, 1008–1013 (2004).

57. Vierkotten, J., Dildrop, R., Peters, T., Wang, B. & Rüther, U. Ftm is a novel basal body protein of cilia involved in Shh signalling. Development 134, 2569–2577 (2007).

58. Dani, A., Huang, B., Bergan, J., Dulac, C. & Zhuang, X. Superresolution imaging of chemical synapses in the brain. Neuron 68, 843–856 (2010).

59. Kanchanawong, P. et al. Nanoscale architecture of integrin-based cell adhesions. Nature 468, 580–584 (2010).

60. Szymborska, A. et al. Nuclear pore scaffold structure analyzed by super-resolution microscopy and particle averaging. Science 341, 655–658 (2013).

61. Huang, B., Jones, S. A., Brandenburg, B. & Zhuang, X. Whole-cell 3D STORM reveals interactions between cellular structures with nanometer-scale resolution. Nat Methods 5, 1047–1052 (2008).

62. Jones, S. A., Shim, S.-H., He, J. & Zhuang, X. Fast, three-dimensional super-resolution imaging of live cells. Nat Methods 8, 499–508 (2011).

63. Sayer, J. A. et al. The centrosomal protein nephrocystin-6 is mutated in Joubert syndrome and activates transcription factor ATF4. Nat Genet 38, 674–681 (2006).

64. Helou, J. et al. Mutation analysis of NPHP6/CEP290 in patients with Joubert syndrome and Senior-Løken syndrome. J Med Genet 44, 657–663 (2007).

65. Baala, L. et al. Pleiotropic effects of CEP290 (NPHP6) mutations extend to Meckel syndrome. Am J Hum Genet 81, 170–179 (2007).

66. Yee, L. E. et al. Conserved Genetic Interactions between Ciliopathy Complexes Cooperatively Support Ciliogenesis and Ciliary Signaling. PLoS Genet. 11, e1005627 (2015).

67. Goetz, S. C. & Anderson, K. V. The primary cilium: a signalling centre during vertebrate development. Nat. Rev. Genet. 11, 331–344 (2010).

68. Byrne, E. F. X. et al. Structural basis of Smoothened regulation by its extracellular domains. Nature 535, 517–522 (2016).

69. Dafinger, C. et al. Mutations in KIF7 link Joubert syndrome with Sonic Hedgehog signaling and microtubule dynamics. J. Clin. Invest. 121, 2662–2667 (2011).

70. Hynes, A. M. et al. Murine Joubert syndrome reveals Hedgehog signaling defects as a potential therapeutic target for nephronophthisis. Proc Natl Acad Sci USA 111, 9893–9898 (2014).

71. You, Y. & Brody, S. L. Culture and differentiation of mouse tracheal epithelial cells. Methods Mol Biol 945, 123–143 (2013).

72. Bossi, M. et al. Multicolor far-field fluorescence nanoscopy through isolated detection of distinct molecular species. Nano Lett. 8, 2463–2468 (2008).

73. Fienup, J. R., Guizar-Sicairos, M. & Thurman, S. T. Efficient subpixel image registration algorithms. Opt. Lett., OL 33, 156–158 (2008).

74. Schneider, C. A., Rasband, W. S. & Eliceiri, K. W. NIH Image to ImageJ: 25 years of image analysis. Nat Methods 9, 671–675 (2012).

75. Pieper, U. et al. ModBase, a database of annotated comparative protein structure models and associated resources. Nucleic Acids Res. 42, D336–46 (2014).

76. Webb, B. & Sali, A. Comparative Protein Structure Modeling Using MODELLER. Curr Protoc Bioinformatics 54, 5.6.1–5.6.37 (2016).

77. Tsirigos, K. D., Peters, C., Shu, N., Käll, L. & Elofsson, A. The TOPCONS web server for consensus prediction of membrane protein topology and signal peptides. Nucleic Acids Res. 43, W401–7 (2015).

78. Drozdetskiy, A., Cole, C., Procter, J. & Barton, G. J. JPred4: a protein secondary structure prediction server. Nucleic Acids Res. 43, W389–94 (2015).

